# *Vibrio cholerae* VgrG1-ACD Targets *E. coli* MreB and Regulates Cellular Morphology

**DOI:** 10.64898/2026.08.07.743456

**Authors:** Amaresh Jana, Amrita Maity, Shubham Das, Saikat Das, Priyanka Dutta, Sankar Maiti

## Abstract

*Vibrio cholerae* employs the Type VI Secretion System (T6SS) to outcompete neighboring bacteria and establish successful colonization within the polymicrobial gut. The T6SS effector VgrG1 contains a C-terminal Actin Crosslinking Domain (VgrG1-ACD), well known for irreversibly crosslinking eukaryotic actin to disrupt the host cytoskeleton. However, whether VgrG1-ACD also contributes directly to interbacterial competition has remained unexplored. Here, we identify a previously unrecognized bacterial target of VgrG1-ACD: MreB, the bacterial actin homolog essential for cell shape, cell wall synthesis, and cytoskeletal organization. We demonstrate that VgrG1-ACD specifically binds *Escherichia coli* MreB, resulting in pronounced morphological defects and impaired bacterial growth. Notably, unlike its activity toward mammalian actin, VgrG1-ACD targets MreB without crosslinking it, revealing a distinct mode of action. These findings expand the functional repertoire of VgrG1-ACD beyond host-directed virulence and uncover a dual-targeting strategy through which a single T6SS effector manipulates both mammalian actin and its bacterial homolog, likely enhancing *V. cholerae* fitness during interbacterial competition and host colonization.

## 1. Introduction

*Vibrio cholerae* is one of the most dangerous waterborne pathogens responsible for enteric disease worldwide. According to the World Health Organization (WHO), the global burden of cholera continued to rise in 2024, with both disease incidence and mortality increasing compared with the previous year^1^. Reported cholera cases increased by 5%, while deaths rose by 50%, resulting in more than 6,000 fatalities worldwide^2^. From January 1 to November 24, 2024, 733,956 cholera cases and 5,162 deaths were reported across 33 countries. India accounted for 10,615 of these cases and 52 associated deaths, underscoring the ongoing global health burden of *Vibrio cholerae* and the critical need for a deeper understanding of the molecular mechanisms underlying its pathogenicity and ecological competitiveness^1^.

More than 200 serotypes of *V. cholerae* exist in the environment, broadly classified by surface O-antigen into the O1/O139 and non-O1/O139 (NOVC) groups^3,4^. O1/O139 strains drive cholera pandemics worldwide, owing to their possession of cholera toxin (CTX) and the toxin-coregulated pilus, which together establish lethal intestinal infection^5–7^. Apart from this CTX-mediated pathogenesis, *V. cholerae* has a sophisticated nanomachine called the Type Six Secretion System (T6SS), which is used for contact-dependent killing of host cells^8^. T6SS consists of 13 core components (TssA-TssM). A TssAEFGK complex baseplate anchors to the membrane via TssJLM complex, supporting an HCP (TssD) tail tube and VgrG (TssI) spike enclosed by a TssBC sheath^9–12^. It functions like a syringe to inject effectors, with ATPase ClpV mediating sheath recycling^10^. Along with targeting eukaryotic system, T6SS is involved in interbacterial competition, and they can potentially outcompete gram-negative bacteria present in the same environmental milieu. Species like *Pseudomonas aeruginosa, Burkholderia thailandensis, Yersinia pseudotuberculosis* and *Vibrio cholerae* employ their T6SS for niche occupancy, scavenging micronutrients like iron, manganese, zinc, and spreading infection^13–15^. Some dominant gram-negative bacterial species like *V. choleare, V. anguillarum, S. enterica serovar Typhimurium, P. aeruginosa, P. syringe* use their T6SS for outcompeting their neighboring cells, present in same environmental milieu^10,16–19^. *Pseudomonas syringe* employs T6SS-1 effectors Tde1 and Tde4 to kill *E. coli* by DNA degradation and changing cellular morphology^20^. Previous study showed, the *V. cholerae* T6SS effector TseH degrades peptidoglycan, resulting in cell wall permeabilization and lysis of *Escherichia coli* cells^21^.

This competition is not restricted to environmental habitats but is equally critical within the gut microenvironment, where a complex community of commensal microbes collectively maintains host health and provides colonization resistance against invading pathogens. Consumption of contaminated food or water can disrupt this microbial balance, leading to gut dysbiosis and creating opportunities for pathogenic enteric bacteria to establish infection^22,23^. Recent studies suggest that interbacterial competition is not merely a defensive strategy but a prerequisite for successful colonization^24^. To access epithelial niches and persist within the host, pathogens must actively eliminate or displace resident microbiota, making interspecies competition a key determinant of infection and disease progression^25,26^. Enteric pathogen *Salmonella enterica* serovar Typhimurium kills commensal *Klebsiella oxytoca* in vitro and also in the mouse gut using Tde4, a T6SS effector^27^. Non-toxigenic *Bacteroides fragilis* protects us from the colonization of enterotoxigenic *B. fragilis* by functional T6SS^28^. *V. cholerae* uses T6SS to increase contraction of the zebrafish intestine and expel the symbiotic *Aeromonas veronii* from the gut rather than direct killing^29^. Enteric pathogen *Citrobacter rodentium* uses T6SS-1 to kill *E. coli* Mt1B1 in murine gut^30^. This raises a fundamental question: what molecular strategies enable pathogens such as *Vibrio cholerae* to outcompete neighbouring bacteria and establish dominance within the intestinal niche? Scientists have demonstrated that the RtxA toxin of *V. cholerae* exhibits potent cytotoxicity against mammalian epithelial cells (Hep-2), characterized by pronounced cell rounding and apoptosis^31,32^. This toxicity is driven by the Actin Crosslinking Domain (ACD), which enzymatically catalyzes the formation of an intermolecular isopeptide bond between the γ-carboxyl group of E270 and the ε-amino group of K50 in an ATP-dependent manner^33^. Subsequent research identified that the *V. cholerae* (classical biotype) VgrG-1 effector similarly harbors a C-terminal ACD (732-1163aa), which is functionally responsible for actin crosslinking both in vitro and in vivo^34^. A previous study from our group has further elucidated that the VgrG-1 ACD contains a specific Actin Binding Motif (ABM) that mediates direct interaction with actin monomers^35^. Furthermore, we have deciphered that T6SS tube protein HCP has been shown to interact directly with actin and colocalize with actin stress fibers upon expression in HeLa cells, highlighting a multifaceted strategy for host cytoskeletal regulation^36^. Likewise, bacterial cytoskeletal proteins have emerged as important targets during interbacterial competition. For example, the chromosomally encoded toxins YeeV, CptA, and CbtA produced by *Escherichia coli* interfere with the polymerization of the essential cytoskeletal proteins MreB and FtsZ, leading to defects in cell division and pronounced changes in bacterial cell morphology, and ultimately causing cell death^37–39^.

These observations suggest that, similar to the eukaryotic cytoskeleton, bacterial cytoskeletal proteins represent vulnerable targets during interbacterial antagonism. Based on this, we hypothesize that the *V. cholerae* VgrG1-ACD may extend its activity beyond eukaryotic targets and instead modulate the dynamics of the bacterial actin homolog, MreB, thereby targeting the cytoskeletal machinery of competing bacteria during interbacterial competition and gut colonization.

In this study, we identify the bacterial actin homolog MreB as a direct target of the *Vibrio cholerae* actin cross-linking domain (ACD_VC_). We show that ACD_VC_ binds to MreB, compromises bacterial cell viability, and perturbs MreB-dependent cell morphology, thereby revealing a previously unappreciated novel function of ACD_VC_ in bacterial cytoskeletal regulation. ACD_VC_ acts as a cytoskeleton-targeting effector during interbacterial competition, substantially expanding its functional repertoire beyond its well-established role in manipulating the mammalian actin cytoskeleton. Together, these findings provide the first evidence that ACD_VC_ can function as a cytoskeleton-targeting effector in interbacterial competition, beyond its established role in targeting the mammalian cytoskeleton.

## 2. Results

### 2.1. Evolutionary and Structural Relationship Between MreB and Rabbit Actin

MreB is a highly conserved bacterial cytoskeletal protein that plays a central role in maintaining the morphology and structural integrity of rod-shaped bacteria^40^. Depletion of MreB results in a loss of rod shape and the adoption of a spherical morphology, underscoring its essential function in bacterial cell shape determination^41^. Beyond its structural role, MreB participates in chromosome segregation and coordinates peptidoglycan biosynthesis, thereby contributing to cell growth and division^42,43^. Similar to rabbit actin, MreB undergoes ATP-dependent polymerization to form dynamic filaments, although the molecular mechanisms governing its assembly and dynamics remain incompletely understood^44,45^. Despite these functional similarities, the two proteins differ substantially in their higher-order filament architecture: actin polymerizes into right-handed, polar double-helical filaments, whereas MreB assembles into straight, antiparallel double filaments^46–48^.

Although previous studies have identified MreB as a structural homolog of mammalian actin a comprehensive comparison integrating evolutionary, sequence, and structural analyses has been lacking^49^. To establish the molecular basis for investigating MreB as a potential target of the ACD_VC_, we performed an integrated comparative analysis of these two cytoskeletal proteins. We first assessed their evolutionary relationship using phylogenetic reconstruction, followed by sequence conservation analysis and structural comparisons of their domain organization and three-dimensional architectures.

A phylogenetic tree constructed using representative bacterial actin homologs and eukaryotic actins revealed a clear evolutionary separation between MreB and eukaryotic actin (Figure 1A). Escherichia coli MreB clustered within a well-defined bacterial MreB clade and exhibited the closest evolutionary relationship to homologs from *Vibrio cholerae*, *Salmonella enterica* serovar Typhimurium, *Shigella flexneri*, and other members of the Enterobacterales, reflecting the high conservation of MreB among rod-shaped bacteria. In contrast, vertebrate, plant, and fungal actins formed a distinct, strongly supported clade, clearly separated from their bacterial counterparts. Despite serving analogous functions as ATP-dependent cytoskeletal proteins, the pronounced phylogenetic separation between the MreB and actin clades underscores their extensive evolutionary divergence (Figure 1A).

**Figure 1:**
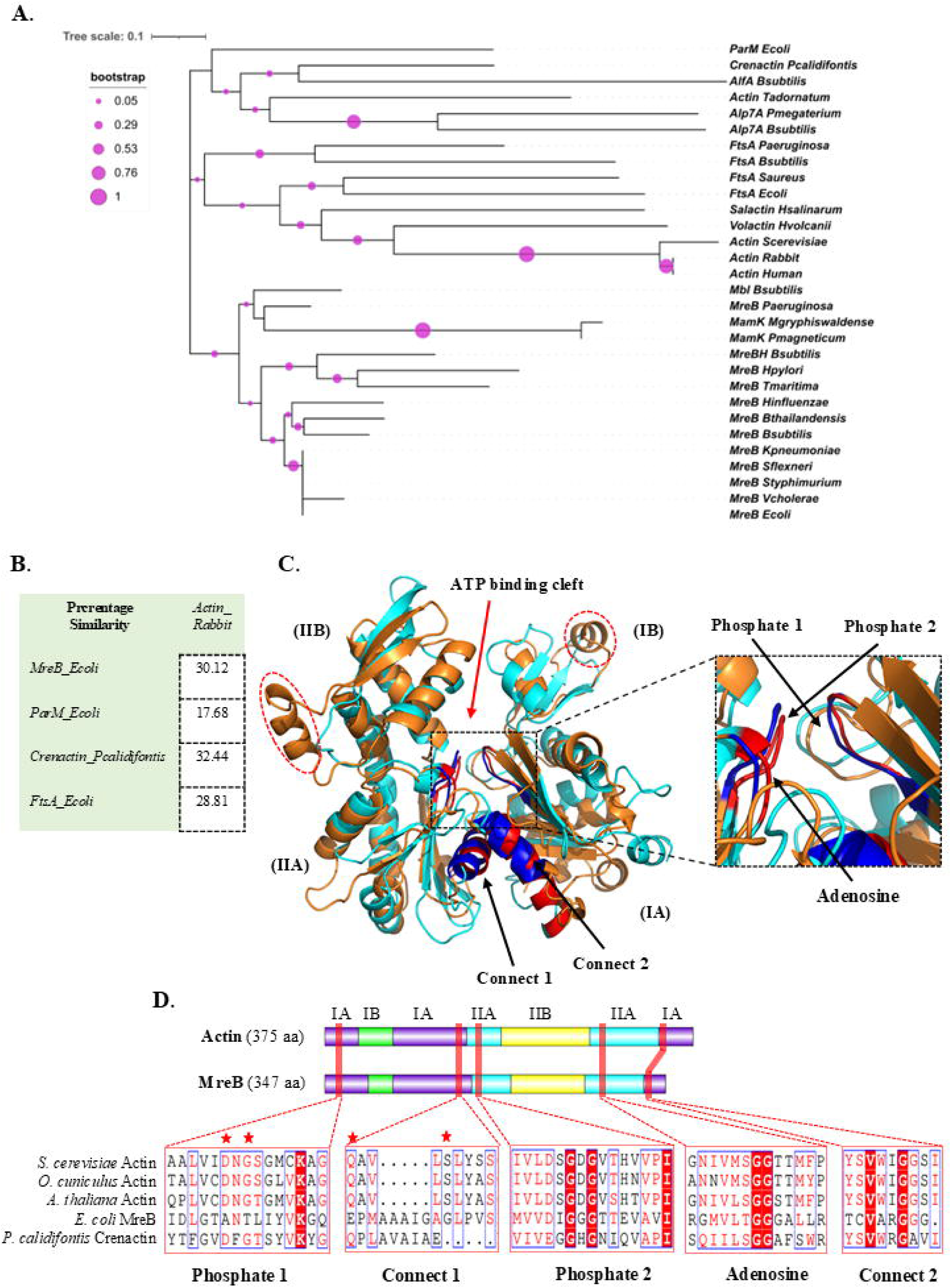
*E. coli* MreB is sequentially distant but structurally homologous to mammalian actin. **(A)** Phylogenetic tree of MreB and and other actin super-family proteins across different kingdom (bacteria, archaea, yeast, plant and mammals). Tree was constructed using Maximum Likelihood method with Gamma-distributed rate variation among sited (G) and 1000 bootstrap replicates. Tree indicated that MreB and actin are evolutionarily related at the level of an ancient common ancestor, but they are not closely related in recent evolutionary terms. **(B)** Structural alignment of actin (orange) and MreB (cyan) showed the overall conservation of different actin folds and ATP binding cleft in MreB despite of low sequence identity. Four Subdomains (IA, IB, IIA and IIB) are marked. Phosphate 1, phosphate 2, adenosine, connect 1, and connect 2 regions in both actin and MreB were highlighted in blue and red respectively. **(C)** Comparative analysis of the overall amino acid sequence similarity between rabbit actin and actin homologs. The percentage similarity values are shown for *E. coli* MreB, *E. coli* ParM, *P. calidifontis* crenactin, and *E. coli* FtsA. (**D)** Schematic representation of the ATP-binding cleft in rabbit actin and bacterial actin homologs. The ATP-binding regions (Phosphate 1, Connect 1, Phosphate 2, Adenosine, and Connect 2), indicated by red boxes, were identified by multiple sequence alignment and comparison with previously reported ATP-binding motifs. Conserved amino acid residues within each ATP-binding region are highlighted by white letters with red background in the sequence alignment. Substituted amino acid positions are denoted in red stars.

Sequence comparison further supported this evolutionary divergence. *E. coli* MreB shared only 15.31% sequence identity with rabbit actin, while FtsA and ParM showed even lower identities of 13.76% and 12.88%, respectively (Figure S1A). Despite this low identity, MreB exhibited the highest sequence similarity to rabbit actin (30.12%), compared with FtsA (28.81%) and ParM (17.68%) (Figure 1B). This suggests that although the amino acid sequences have diverged, MreB has retained many conserved residues that help maintain the characteristic actin fold and ATP-dependent function. This prompted us to further investigate the structural relationship between MreB and actin through three-dimensional structural alignment and domain organization analysis. We first generated a high-confidence (pTM score 0.93) structural model of *E. coli* MreB using AlphaFold and compared it with the previously determined crystal structure of *Thermotoga maritima* MreB. Sequence alignment revealed 53.43% sequence identity and 64.61% sequence similarity between the two proteins (Figure S1A). Additionally, structural superposition between *E. coli* and *T. maritima* MreBs yielded an RMSD of 0.66 Å, indicating that our predicted *E. coli* MreB model is highly consistent with the experimentally resolved MreB structure (Figure 1C). We next compared the *E. coli* MreB structure with representative members of the actin superfamily. Structural superposition showed that MreB aligned well with rabbit actin, exhibiting an RMSD of 4.4 Å over 255 equivalent residues (Figure 1C, S1C). A similar degree of structural conservation was observed with ParM (RMSD = 4.3 Å over 247 equivalent residues), whereas FtsA displayed substantially greater divergence (RMSD = 14.3 Å over 232 equivalent residues) (Figure S1B, C). Despite the overall conservation of the actin fold, MreB lacked two α-helices located on the outer surface of actin subdomains IB and IIB, representing a prominent structural distinction between the two proteins (Figure 1C).

Domain organization analysis further revealed a highly conserved ATP-binding core despite differences in overall protein length (Figure 1D, S2). Both MreB and rabbit actin possess the characteristic actin fold comprising subdomains IA, IB, IIA, and IIB (Figure 1C, S2)^49,50^. Multiple sequence alignment of the nucleotide-binding pocket demonstrated strong conservation of the phosphate 1 (Asp <u>x</u> Gly), connect 1 (Gln <u>x</u> <u>x</u> <u>x</u> Ser), phosphate 2 (Asp <u>x</u> Gly <u>x</u> Gly), adenosine (Gly Gly), and connect 2 (Gly) motifs between *E. coli* MreB and actins from *Saccharomyces cerevisiae*, *Oryctolagus cuniculus*, and *Arabidopsis thaliana* (Figure 1D)^51^. Key residues involved in ATP binding and hydrolysis were largely conserved, with only a few substitutions observed within the phosphate 1 and connect 1 motif (Figure 1D). In *E. coli* MreB, the conserved Asp residue in the phosphate 1 motif was replaced by Ala20 (denoted with red star), representing a non-conservative substitution that removes a negatively charged side chain, while the conserved Gly was substituted by Thr22 (denoted with red star), introducing a polar residue in place of a small, flexible amino acid. In the connect 1 motif, Gln was conservatively replaced by Glu143 (denoted with red star), whereas Ser was substituted by Gly134 (denoted with red star), a change that is expected to increase local flexibility while preserving the overall architecture of the nucleotide-binding pocket (Figure 1D).

Collectively, our findings support the previously established notion that MreB is a structural homolog of eukaryotic actin, sharing the conserved actin fold and ATP-binding architecture. Beyond this, our study uncovers characteristic amino acid substitutions in the Phosphate 1 and Connect 1 motifs, as well as structural remodelling of subdomains IB and IIB, underscoring previously underappreciated structural adaptations unique to bacterial MreB.

### 2.2. In Silico Study Showed *Vibrio cholerae* Actin Crosslinking Domain and *E. coli* MreB Forms a Stable Complex

Many bacterial pathogens manipulate the mammalian actin cytoskeleton to promote infection, intracellular motility, and cell-to-cell spread through direct interaction with actin or by targeting actin-binding proteins^52^. In *Vibrio cholerae*, the Type VI Secretion System (T6SS) effector VgrG1 contains an actin cross-linking domain (ACD_VC_) that covalently cross-links mammalian actin and disrupts host cytoskeletal dynamics^33^. Based on the structural and functional conservation between mammalian actin and the bacterial actin homolog MreB, we hypothesized that MreB could represent a previously unrecognized target of ACD_VC_ during T6SS-mediated interbacterial competition. If so, ACD_VC_-mediated perturbation of MreB in neighboring bacterial cells could impair cytoskeletal function and provide *V. cholerae* with a competitive advantage within polymicrobial environments.

To gain insights into the direct interaction between *E. coli* MreB and the ACD_VC_, we performed molecular docking studies. The three-dimensional (3D) structures of both MreB (347 amino acids) (pTM score 0.93) and VgrG1-ACD (residues 732-1163) (pTM score 0.79) were predicted using AlphaFold Colab (Figure 2A, B). Previously resolved crystal structure of VgrG1-ACD (PDB: 4DTD) lacks the C-terminal region (residues 1080-1163)^53^. Therefore, we predicted the full-length ACD_VC_ structure (residues 732-1163) for this study. Superposition of the predicted model with the crystal structure, excluding the unresolved C-terminal region, yielded an RMSD of 1.068 Å over 2,372 aligned atoms, indicating that the AlphaFold model closely resembles with the experimentally resolved structure (Figure S3A). The quality of the predicted models was assessed using the Predicted Aligned Error (PAE) heatmap^54^. The PAE heatmap provides a two-dimensional representation of the expected positional error (in Å) between pairs of residues, enabling evaluation of the confidence in the relative positioning of different regions within the predicted structure. In the heatmap, blue regions indicate high-confidence predictions with low expected error, whereas red regions represent lower-confidence predictions with higher expected error. For MreB, the PAE heatmap was predominantly blue, indicating a high level of confidence across the predicted structure (Figure 2B). Likewise, the ACD_VC_ model exhibited an overall high-confidence prediction, with most regions displaying low predicted alignment error. However, a segment spanning residues 350-430 of the predicted ACD_VC_ model (corresponding to residues 1080-1163 of VgrG1) exhibited relatively lower confidence than the rest of the structure (Figure 2A). Although this C-terminal region appeared internally consistent, suggesting that it adopts a structured fold, its orientation relative to the main domain remained uncertain, indicating increased conformational flexibility and reduced confidence in its relative positioning. These predictions were further supported by their secondary structural orientation obtained from PSIPRED (Figure S3B, C).

**Figure 2.**
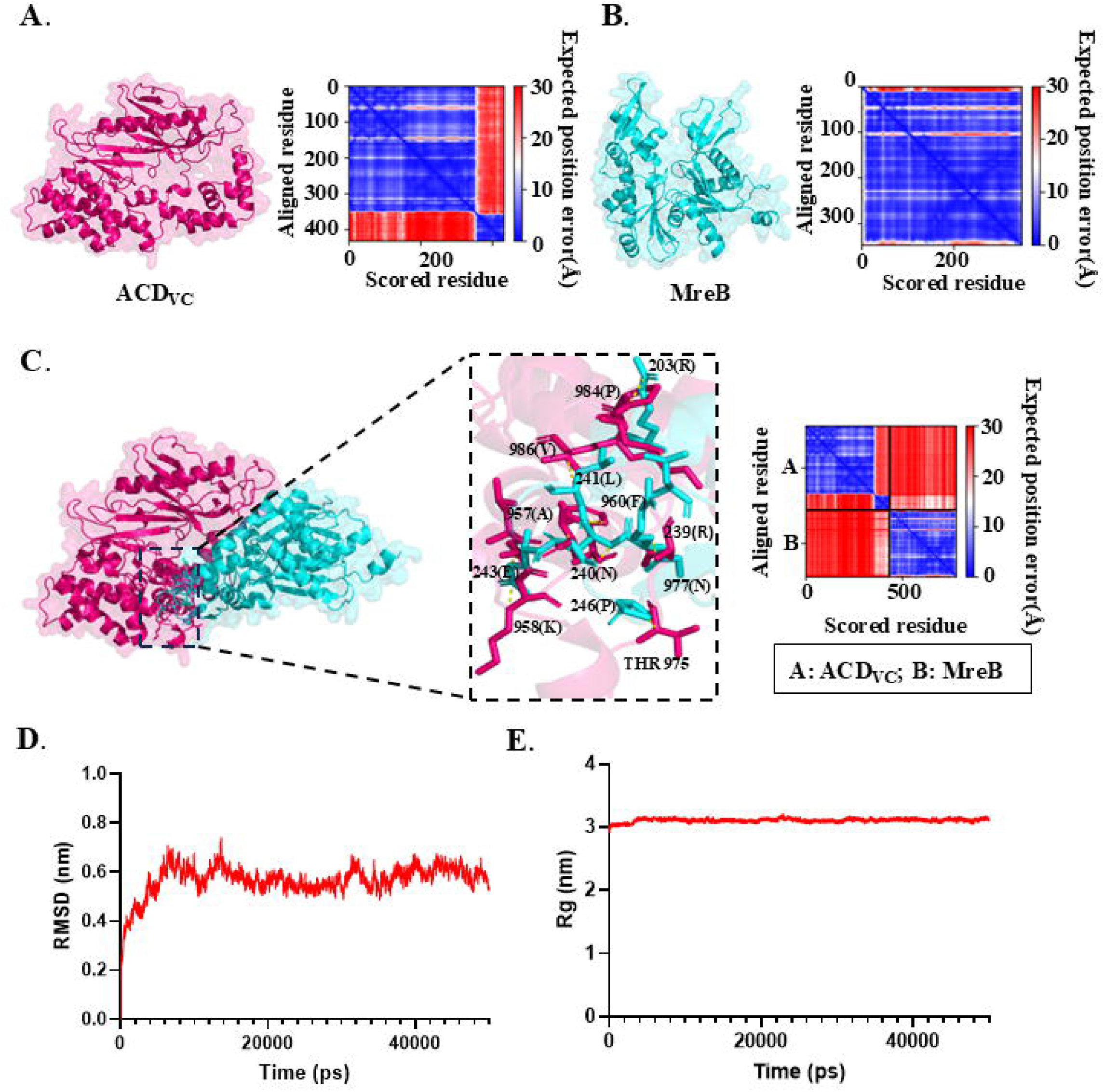
Stable ACD_VC_-MreB complex predicted by docking and MD simulations. Predicted three-dimensional structures of **(A)** the VgrG1 Actin cross-linking domain (ACD; residues 732-1163; pink) and **(B)** *E. coli* MreB (cyan) generated using AlphaFold2. The predicted aligned error (PAE) plots displayed adjacent to each model indicate the confidence of residue-residue positional relationships within the predicted structures. Where blue tiles represent high confidence and red for low confidence in predicted structure. **(C)** Predicted 3D model of the MreB-ACD_VC_ complex generated using AlphaFold2-Multimer. The corresponding predicted aligned error (PAE) plot is shown alongside the complex model, while the enlarged view depicts the predicted interaction interface, highlighting ACD_VC_ residues (pink; Ala957, Lys958, Phe960, Thr975, Asn977, Pro984, and Val986) and MreB residues (cyan; Arg203, Arg239, Asn240, Leu241, Glu243, and Pro246) involved in the predicted binding contacts. **(D)** Plot showing the backbone root mean square deviation (RMSD) of the ACD-MreB complex relative to the initial structure throughout the simulation. **(E)** Radius of gyration (Rg) plot depicting the structural compactness and overall stability of the ACD-MreB complex during the 50 ns simulation.

Having obtained high-confidence structural models, we next investigated the interaction between MreB and ACD_VC_ through in silico protein-protein docking using AlphaFold Colab-Multimer (Figure 2C). Analysis of the highest-ranked docked complex revealed a putative interaction interface predominantly stabilized by hydrogen bonds. On the ACD_VC_, residues Ala957, Lys958, Phe960, Thr975, Asn977, Pro984, and Val986 formed interactions with Arg203, Arg239, Asn240, Leu241, Glu243, and Pro246 of MreB (Figure 2C). Notably, these interacting MreB residues are located within subdomain IIB, which is structurally homologous to subdomain 4 of the rabbit actin.

To further evaluate the stability and dynamic behavior of the predicted MreB-ACD_VC_ complex, a 50 ns molecular dynamics (MD) simulation was performed using GROMACS (Video S1)^55^. The root mean square deviation (RMSD) of the complex remained below 0.7 nm throughout the simulation, indicating that the complex maintained its overall structural integrity relative to the initial conformation (Figure 2D). Additionally, the radius of gyration (Rg), which reflects the compactness of the complex, remained stable over the 50 ns trajectory, with an average Rg value of 3.11 nm (component values: Rgx = 2.77 nm, Rgy = 2.09 nm, and Rgz = 2.70 nm) (Figure 2E). Together, these findings indicate that the predicted MreB-ACD_VC_ complex retained its structural stability and overall compactness throughout the simulation.

### 2.3. Biochemical assays Reveals Direct Association of ACD_VC_ with *E. coli* MreB

To investigate the direct interaction between ACD_VC_ and MreB, they were cloned and expressed as N-terminal 6X-His tagged proteins in *Escherichia coli* BL21(DE3) cells and purified^56^. SDS-PAGE analysis demonstrated high-purity protein preparations, with major bands migrating at the expected molecular masses of approximately 40 kDa for MreB and 51 kDa for ACD_VC_, respectively (Figure 3A, B).

**Figure 3:**
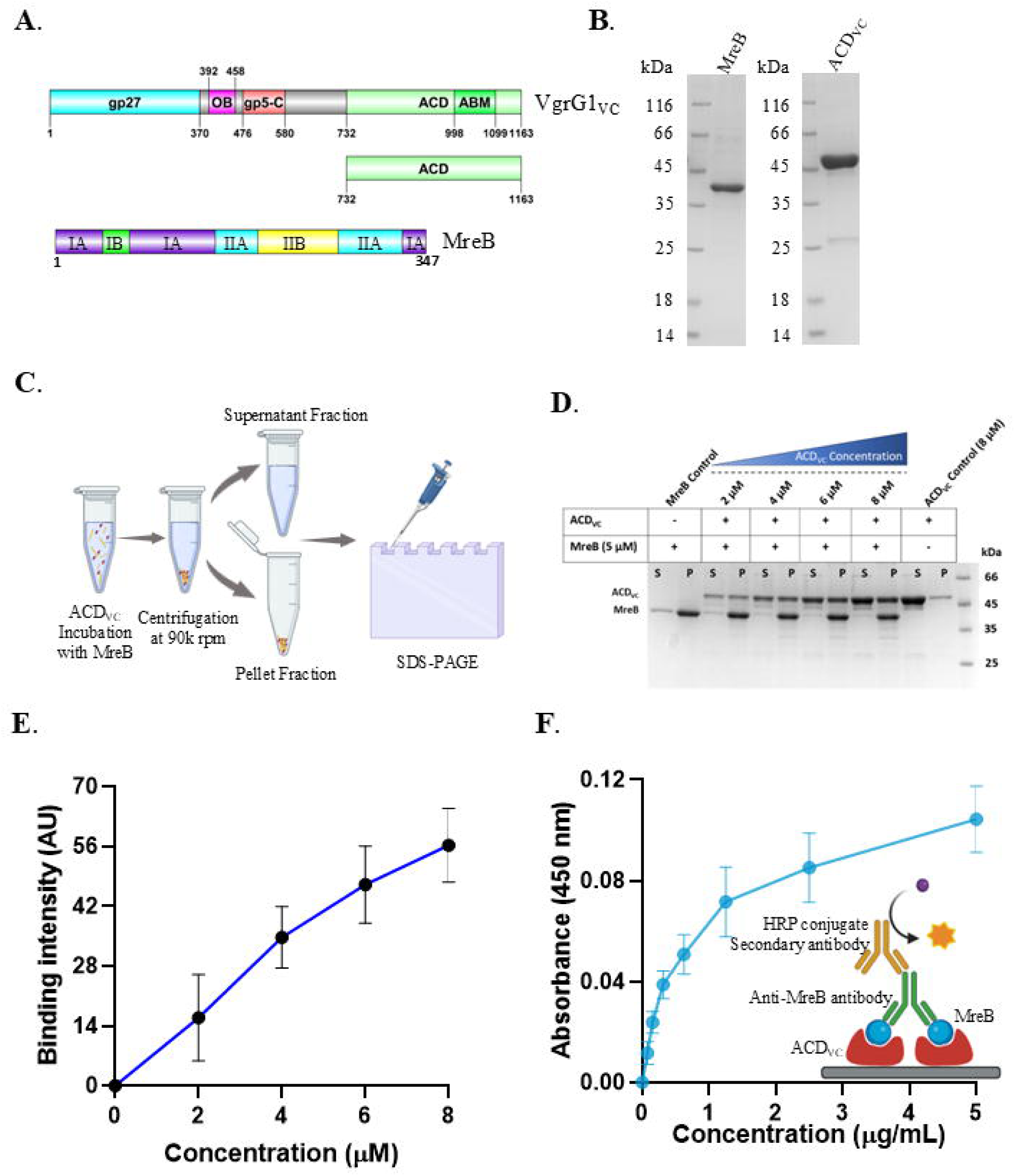
ACD_VC_ directly interacts with MreB. **(A)** Schematic representation of different domains in VgrG1_VC_ (1163 aa) and MreB (347 aa). **(B)** Coomassie-stained 10% SDS-PAGE gel showing purified 6X-His tagged-MreB (347 aa) and ACD_VC_ (732-1099 aa) proteins **(C)** Schematic experimental settings of high-speed MreB co-sedimentation assay: ACD_VC_ incubated with MreB_EC_ in room temperature for 30 mins followed by centrifugation at 90K. Supernatant (S) and pellet (P) collected separately and analyzed by SDS-PAGE. **(D, E)** Concentration-dependent interaction between MreB and ACD_VC_. Increasing concentrations of ACD_VC_ resulted in a progressive increase in the MreB-bound fraction, as shown by the representative gel image and corresponding densitometric analysis. Quantification of the bound ACD_VC_ demonstrated a concentration-dependent increase in MreB-ACD_VC_ interaction. **(F)** ELISA-based analysis of the MreB-ACD_VC_ interaction. The schematic illustrates the assay principle, in which immobilized ACD_VC_ captures MreB, followed by detection using anti-MreB antibody and an HRP-conjugated secondary antibody. The binding curve shows a concentration-dependent increase in absorbance at 450 nm, indicating specific interaction between ACD_VC_ and MreB.

Then, we performed the high-speed co-sedimentation assay to examine the interaction between MreB and ACD_VC_. MreB was polymerised in buffer containing, 5 mM Tris-HCl (pH 7.5), 0.1 mM CaClL, 0.2 mM ATP, and 5 mM MgCl_2_. Increasing concentrations of purified ACD_VC_ were incubated with *E. coli* MreB in the previously described polymerization buffer at 25°C for 30 min. MreB alone served as a positive control for polymerization, whereas ACD_VC_ alone was used to confirm the solubility of the protein. Following high-speed centrifugation, the supernatant and pellet fractions were run in 10% SDS-PAGE to assess MreB-ACD_VC_ association (Figure 3C). In the test samples, ACD_VC_ recovered in the pellet fraction along with MreB in an increasing concentration dependant manner. Whereas, in absence of MreB, ACD_VC_ remained predominantly in the supernatant. (Figure 3D). Further, we had measured the band intensity of ACD_VC_, present in the pellet fraction. And, it showed a concentration-dependent increase in pellet-associated ACD_VC_ (Figure 3E). Together, the results indicate that ACD_VC_ specifically associates and co-sediments with the polymerized MreB.

Furthermore, this interaction was validated by ELISA. A polyclonal antibody against MreB was generated by immunizing BALB/c mice with purified His-tagged *Escherichia coli* MreB. Terminal bleeds were collected, and the specificity of the anti-MreB antisera was confirmed by immunoblot analysis using purified recombinant His-tagged MreB (Figure S5). In ELISA, ACD_VC_ was coated onto the ELISA plate as the ligand, and MreB was used as the analyte. The complex was captured using mouse anti-MreB antibody and detected with HRP-conjugated anti-mouse IgG antibody following the addition of TMB (Figure 3E). Data showed, the absorbance increased in a concentration-dependent manner with increasing concentrations of MreB, whereas no significant change in absorbance was detected in the buffer control (Figure 3E). Along with co-sedimentation, this result further supports that ACD_VC_ interacts with MreB.

### 2.4. ACD_VC_ Does Not Cross-link MreB Despite Its Structural Homology to Actin

*Vibrio cholerae* ACD_VC_ interacts with cellular actin and covalently cross-links actin monomers both in vitro and in-vivo in an ATP-dependent manner^33^. MreB being considered as an actin homolog, we hypothesized that ACD_VC_ could have a role in MreB crosslinking.

For this, purified *E. coli* MreB was incubated with increasing concentrations of ACD_VC_, where actin used as a positive control. After running the samples in SDS-PAGE, we found that ACD_VC_ had significantly crosslinked the actin, and higher molecular weight bands of crosslinked actin were clearly visible (Figure 4A)^57^. Whereas there were no higher molecular weight bands seen in the case of MreB (Figure 4A). These results indicate that ACD_VC_ does not crosslink the bacterial actin homolog MreB. Although a faint high-molecular-weight band was observed in the MreB samples at ∼70 kDa, it does not represent an ACD_VC_-mediated crosslinked product, as the same band was also detected in the MreB-only control and was consistently present throughout protein purification. These additional bands most likely correspond to co-purifying molecular chaperones. In particular, the ∼70 kDa band is consistent with DnaK, the bacterial Hsp70^58^. However, this raised an important question: despite being a structural homolog of actin, why is MreB not crosslinked by ACD_VC_ in the same manner as mammalian actin?

**Figure 4:**
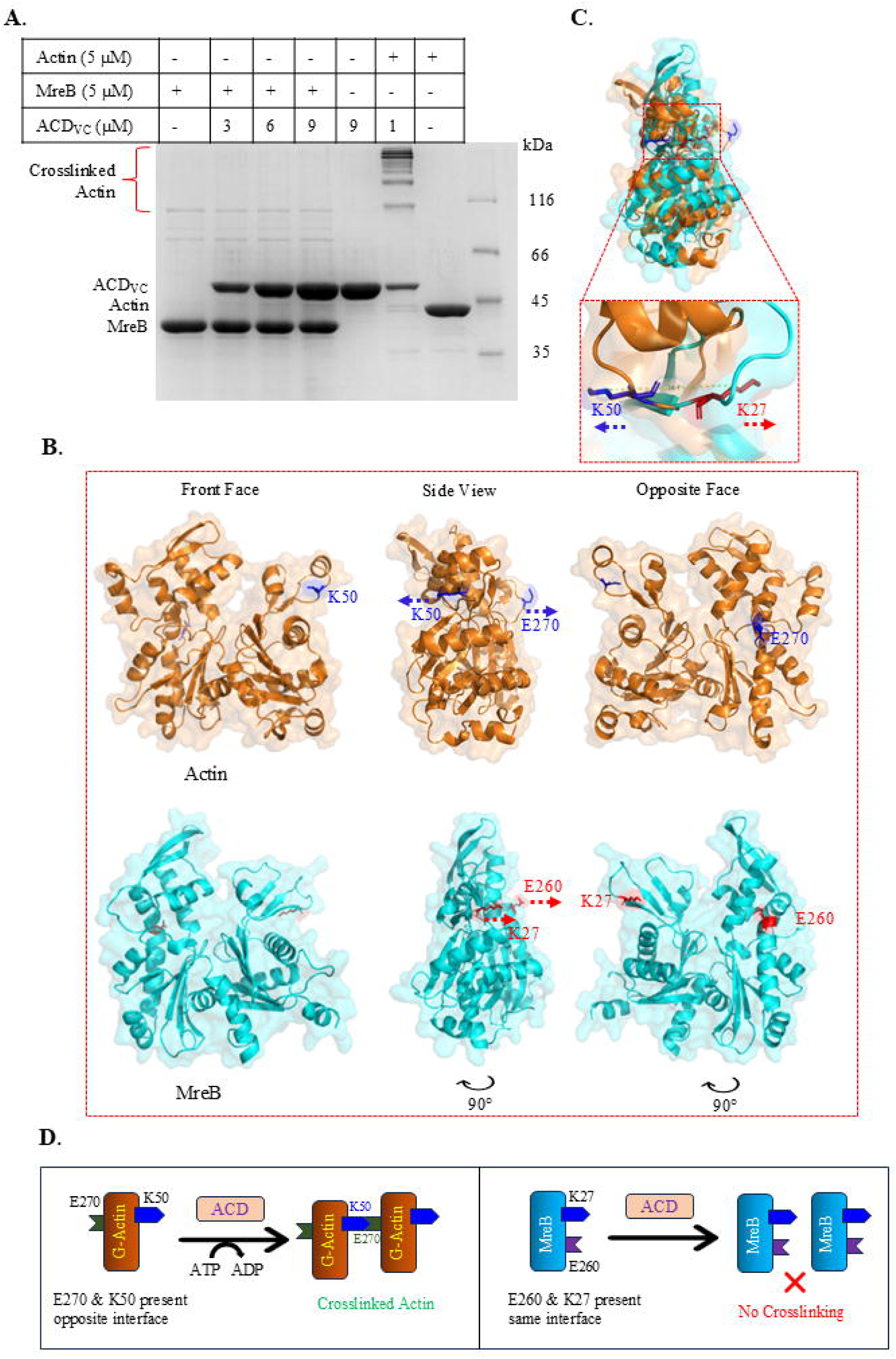
Structural homology is insufficient for ACD_VC_-Mediated crosslinking of MreB. **(A)** Purified MreB (5 µM) was incubated with increasing concentrations of ACD_VC_ (3, 6, and 9 µM), while G-actin (5 µM) was incubated with ACD_VC_ (1 µM) as a positive control. Coomassie-stained 10% SDS-PAGE revealed the formation of high-molecular-weight crosslinked actin species in the presence of ACD_VC_ (lane 6). In contrast, no corresponding crosslinked MreB species were detected (lane 2-4) **(B)** Positional comparison of the residues involved in ACD_VC_-mediated crosslinking, present on actin and MreB. The Lys50 and Glu270 residues of actin (orange; blue labels) and the corresponding Lys27 and Glu260 residues of MreB (cyan; red labels) are shown in front (0°), side (90°), and opposite (180°) orientations to illustrate differences in their surface positioning and spatial arrangement. **(C)** Structural alignment of actin (orange) and MreB (cyan) showing the spatial arrangement of the Lys50 (blue) and Lys27 (red) residues respectively. The magnified view demonstrates that, despite occupying a similar structural position, the Lys side chains are oriented in opposite directions in actin and MreB. **(D)** Proposed model for the substrate specificity of ACD_VC_. In actin (orange), ACD_VC_ promotes the formation of an isopeptide bond between Lys50 (blue) of one G-actin monomer and Glu270 (green) of a second G-actin monomer, leading to actin crosslinking. In contrast, the corresponding residues in MreB (cyan), Lys27 (blue) and Glu260 (purple), are located on the same molecular interface, likely precluding the geometry required for ACD_VC_-mediated crosslinking.

To investigate the reason behind this, we analysed and compared the 3D-structural aspects of actin and MreB. For this, the crystallographic structure of actin (PDB:1J6Z) and high-confidence AlphaFold structure of *E. coli* MreB were considered. During ACD_VC_-mediated crosslinking, an isopeptide bond is formed between K50 of one actin monomer and E270 of a second actin monomer. Here, we observed that in the case of MreB, there was a Lys at the 27^th^ position (at IB subdomain) same as K50 in actin, but no Glu was present at the 270^th^ position (at IIB subdomain) (Figure S2). Further structural alignment of actin and MreB revealed that a Glu was present at the 260^th^ position of MreB, similar to that of actin E270 (Figure 4B). However, K50 and E270 were positioned at opposite surfaces on actin, whereas in MreB K27 and E260 were positioned on the same surface (Figure 4B). Additionally, K50 residues of actin and K27 of MreB occupy similar structural positions but are oriented in opposite directions, forming an angle of approximately 180° with respect to each other (Figure 4C). The distance between the side-chain amino groups of these two Lys residues was approximately 16.7 Å (Figure 4C).

Together, these findings demonstrate that, unlike actin, *E. coli* MreB is not crosslinked by ACD_VC_ (Figure 4D). Structural differences in the spatial arrangement of the substrate residues likely account for the inability of ACD_VC_ to catalyze MreB crosslinking.

### 2.5. ACD_VC_ Adversely Affects the Growth of *E. coli* BL21 (DE3) Cells

To further investigate the implications of MreB-ACD_VC_ interaction on bacterial growth, we performed a bacterial survivability assay in *E. coli* BL21 (DE3) cells.

ACD_VC_ was expressed in BL21 (DE3) cells as a 6X-His-tagged protein, while the pET28a(+) vector served as the vector control. To rule out any growth defect caused simply by protein expression, we used a distantly related 6X-His-tagged mouse Profilin (mPfn) as an additional control. Cultures were serially diluted and spotted onto LB agar plates. At 37°C, ACD_VC_-expressing cells formed colonies up to the 10^-6^ dilution, similar to the induced vector control, mPfn, and all uninduced samples (Figure S6).

In contrast, induction at 40°C markedly reduced the survivability of ACD_VC_-expressing cells. While the induced vector control, mPfn, and all uninduced cultures showed comparable growth up to the 10^-6^ dilution, ACD_VC_-expressing cells exhibited an approximately 100-fold reduction in growth (Figure 5A). To validate this effect, we measured viable cell counts under the same growth conditions. Serially diluted cultures were plated on kanamycin-containing LB agar, and the log CFU/mL was determined. ACD_VC_-expressing cells showed a clear reduction in viable cell counts compared with the vector and mPfn controls. (Figure 5B). Further, the reduction in log CFU/mL count was found to be statistically significant (*P* < 0.05) for ACD_VC_ induced cells with a *P* value of 0.0127 in One-way ANOVA (Figure 5B). Collectively, these findings demonstrate that although ACD_VC_ does not induce MreB crosslinking, it perturbs cellular function and inhibits growth.

**Figure 5:**
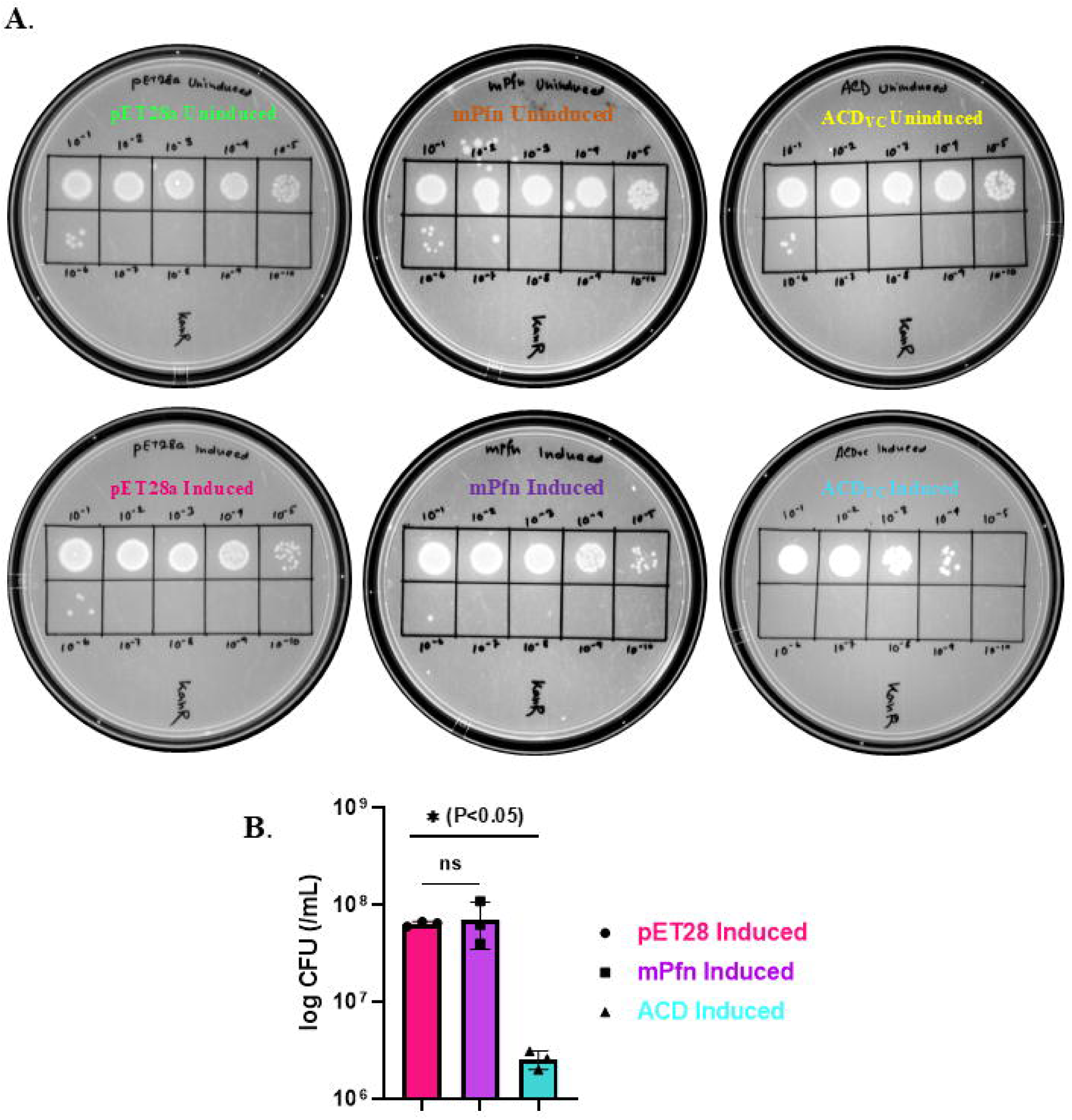
Over-expression of T6SS effector ACD_VC_ inhibits the growth of *E. coli* BL21 (DE3) cells. **(A)** Representative spot assay plates showing the survivability of *E. coli* BL21(DE3) cells expressing ACD_VC_, mouse Profilin (mPfn), or the empty pET28a(+) vector under uninduced and IPTG-induced conditions. Ten-fold serial dilutions (10^-1^-10^-10^) of the cultures were spotted onto kanamycin-containing LB agar plates. A marked reduction in colony formation was observed in ACD_VC_-induced cells compared with the vector and mPfn controls, indicating impaired cell viability following ACD_VC_ expression (*n* = 3 independent experiments). **(B)** Quantification of cell viability following induction of ACD_VC_ expression. Colony-forming units (CFU/mL) are shown for cells expressing the empty pET28a(+) vector (magenta), mouse Profilin (mPfn; purple), and ACD_VC_ (cyan). Data represent the mean ± SD of three independent experiments (n = 3). Statistical significance was determined by one-way ANOVA (*P* < 0.05; ns, not significant).

### 2.6. ACD_VC_ expression Induces Cell Elongation and Bulge Formation

To find out the mechanism behind this growth inhibition, we expressed ACD_VC_ in *E. coli* BL21 (DE3) cells and performed scanning electron microscopy (SEM). Cells harboring the pET28a(+) vector alone served as the negative control. Protein expression was induced at both 37°C and 40°C, after which the cells were fixed with 2.5% glutaraldehyde followed by osmium tetroxide and examined under SEM. Cell lengths were subsequently measured for samples induced at 40°C (Figure 6B). The pET28a(+) vector control exhibited no appreciable change in cell length upon induction, with induced and uninduced cells displaying average lengths of 2.1 ± 0.4 µm and 2.1 ± 0.5 µm (mean ± SD), respectively (Figure 6A, C). Similarly, uninduced ACD_VC_-harboring cells showed an average length of 2.2 ± 0.6 µm, comparable to the vector control. In contrast, induction of ACD_VC_ expression resulted in a marked increase in cell length, with an average length of 2.9 ± 1.5 µm (Figure 6A, C). Notably, a subset of ACD_VC_-expressing cells exhibited pronounced filamentation, reaching lengths of up to 9 µm (Figure 6C). Further, this increase in cell length was found to be statistically significant (P = 0.0005; *P* < 0.05) using One-way ANOVA. It indicated that ACD_VC_ expression induces a filamentous phenotype in *E. coli* (Figure 6C).

**Figure 6:**
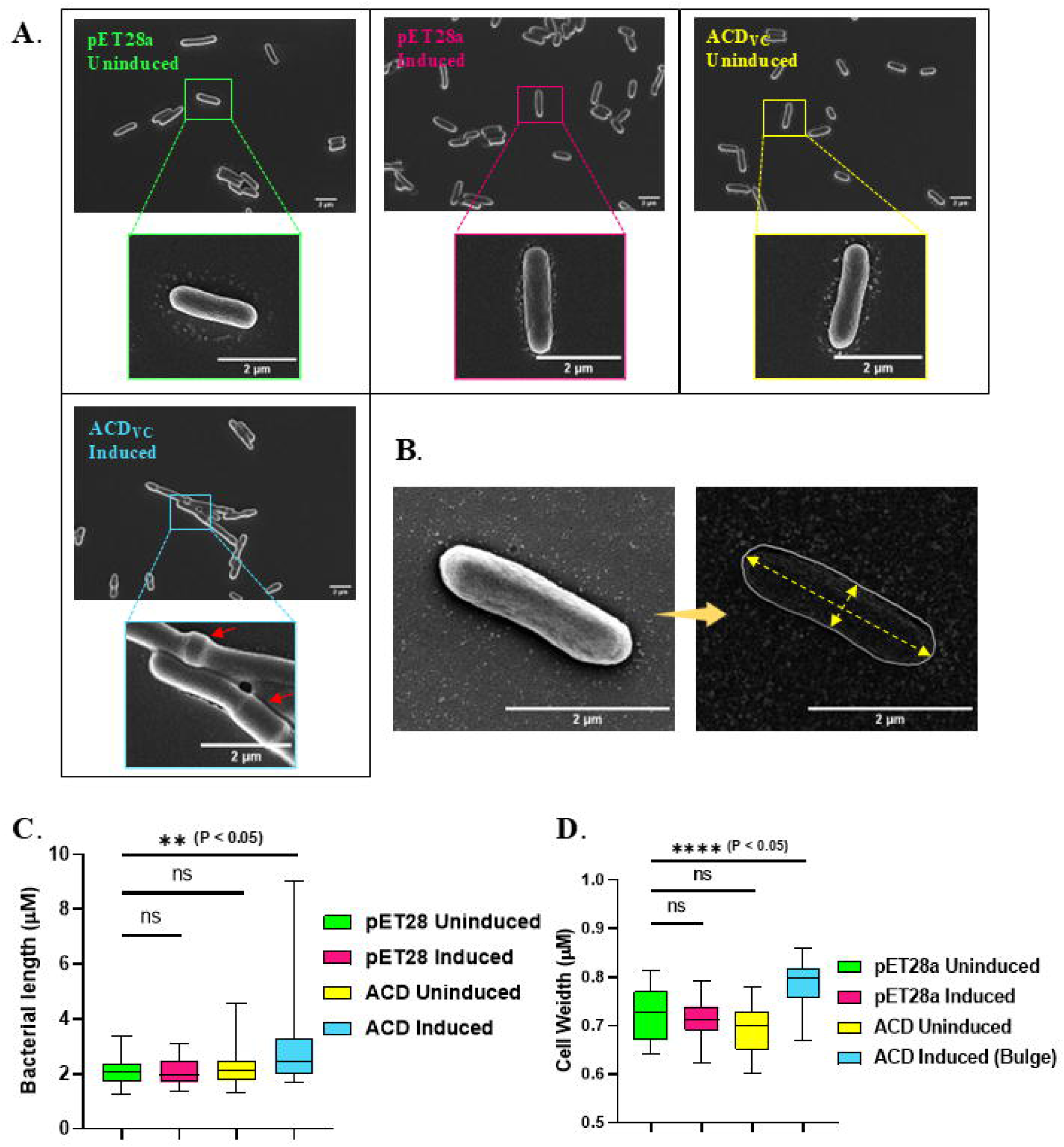
ACD_VC_ expression leads to the morphological changes in *E. coli* BL21 (DE3) cells. **(A)** Representative SEM images of *E. coli* BL21(DE3) cells expressing the empty pET28a(+) vector under uninduced (green) and induced (magenta) conditions, and ACD_VC_ under uninduced (yellow) and induced (cyan) conditions. The lower panels show magnified views of the boxed regions. Cells expressing ACD_VC_ exhibited pronounced morphological abnormalities, including lateral bulge formation (red arrows) and increased cell length. **(B)** Schematic representation of bacterial cell length and width measurements from SEM images. The cell outline indicates the ImageJ-based measurement of length and width (dashed yellow line). **(C-D)** Quantitative analysis of bacterial cell morphology for the empty pET28a(+) vector under uninduced (green) and induced (magenta) conditions, and ACD_VC_ under uninduced (yellow) and induced (cyan) conditions. **(C)** Cell length and **(D)** cell width was measured from SEM images. ACD_VC_-induced cells exhibited a significant increase in both cell length and width, with the latter corresponding to the observed bulged morphology, compared with the control groups. Data represent three independent experiments (n = 33 cells); statistical significance was determined by one-way ANOVA (*P* < 0.05). Scale bar 2 µm.

Additionally, all ACD_VC_-expressing cells exhibited prominent bulges along their lateral cell wall (Figure 6A). Cells with moderate elongation generally displayed a single bulge, whereas progressively elongated cells frequently exhibited multiple bulges, with up to three bulges observed in individual cells. This apparent increase in cell width prompted us to quantitatively assess the morphological changes by measuring the width of cells from each experimental group (Figure 6B). The analysis revealed that ACD_VC_-expressing cells had an average width of 0.80 ± 0.04 µm (mean ± SD), whereas vector-induced, vector-uninduced, and ACD_VC_-uninduced cells exhibited average widths of 0.71 ± 0.03 µm, 0.72 ± 0.05 µm, and 0.69 ± 0.04 µm, respectively (Figure 6A, D). Thus, all control groups displayed comparable cell widths, while ACD_VC_ expression resulted in a noticeable increase in cell width. To determine whether this increase was statistically significant, we performed a one-way ANOVA. The analysis demonstrated a highly significant increase in the width of ACD_VC_ - expressing cells compared with the control groups (*P* < 0.05), indicating that ACD_VC_ expression induces not only cell elongation but also lateral cell expansion in *E. coli* (Figure 6D). A similar morphological phenotype was also observed in cells grown at 37°C (Figure S6). Since MreB is the primary cytoskeletal determinant responsible for maintaining the rod-shaped morphology of bacteria, these observations, together with our biochemical interaction data, strongly support the hypothesis that ACD_VC_ has a clear and significant role in regulating bacterial actin homolog MreB, leading to changes in cell morphology and growth.

## 3. Discussion

In this study, we identify the *Vibrio cholerae* T6SS effector ACD_VC_ as a potential regulator of the bacterial actin homolog MreB. Previous studies established MreB as an actin homolog because it shares key functional properties with eukaryotic actin. Like actin, MreB undergoes ATP-dependent polymerization and plays a central role in maintaining the rod-shaped morphology of bacteria^45^. In addition to controlling cell shape, MreB is involved in cell growth, polarity, protein localization, cell division, chromosome segregation, and DNA replication^42,43^.

To explore whether ACD_VC_ could also target MreB, we first systematically examined the evolutionary and structural relationship between eukaryotic actin and *E. coli* MreB. We began with a phylogenetic analysis of representative actin superfamily proteins from bacteria, archaea, and eukaryotes. We found that actin and MreB positioned into distinct clades corresponding to functional families (Figure 1A). Eukaryotic actins from *Homo sapiens*, rabbit, and *Saccharomyces cerevisiae* clustered tightly together with strong support, consistent with their shared origin and high sequence conservation. Archaea actin homologs crenactin, salactin, and volactin formed neighboring branches, consistent with their close evolutionary relationship to eukaryotic actin (Figure 1A). In contrast, bacterial actin-like proteins exhibited clear functional diversification during evolution. All bacterial MreBs clustered together, suggesting descent from a common ancestral protein. Conversely, FtsA diverged into an independent clade consistent with its role in cytokinesis, while ParM, AlfA, and Alp7A evolved as distinct lineages associated with plasmid segregation and chromosome organization, respectively^59–63^.

Actin homologs have traditionally been classified based on their conserved actin-superfamily fold and functional similarity. Notably, MreB is the closest prokaryotic counterpart of mammalian actin, retaining the canonical fold and a remarkably similar overall size. However, our data showed that *E. coli* MreB shares low sequence identity but relatively high sequence similarity with actin (Figure 1B, S1A). This gap between low identity and comparatively high similarity suggests that MreB has preferentially retained residues important for preserving the actin fold and ATP-dependent activity, even as the primary sequence has diverged extensively. In other words, sequence-level divergence has been tolerated everywhere except where it would compromise the shared molecular framework. This conservation was further supported by structural superposition of rabbit actin and *E. coli* MreB. It showed that MreB was closely aligned with actin (RMSD 4.4 Å over 255 equivalent residues) (Figure 1C). MreB and actin both adopt the canonical actin fold, comprising four subdomains (IA, IB, IIA, IIB) arranged around a conserved ATP-binding core, having a difference in overall protein length (Figure 1C, D, S2). Contrastingly, MreB lacks two surface-exposed α-helices corresponding to actin’s subdomains IB and IIB, indicating that MreB has undergone localized structural adaptation while otherwise preserving the actin-like architecture (Figure 1C). This suggests, both the protein have diverged significantly to perform their specialized functions inside two different cells. Additionally, multiple sequence alignment of the nucleotide-binding pocket revealed strong conservation of the phosphate 2, adenosine, and connect 2 motifs between *E. coli* MreB and actins from *S. cerevisiae*, *Oryctolagus cuniculus*, and *Arabidopsis thaliana*. Only a handful of substitutions were found, in the phosphate 1 and connect 1 motif (Figure 1D). Together, these indicates that, although the amino acid composition of the ATP-binding pocket has diverged over an evolutionary time-scale, its overall structural organization and catalytic framework remain largely conserved. This evolutionary divergence may explain why, dynamic properties of actin homologs, exhibits differences in filament architecture, filament orientation, ATP hydrolysis kinetics, and polymerization behavior^64^. The conservation of these structural features, together with its ATP-dependent polymerization and cytoskeletal functions, supports the view that MreB represents an evolutionary ortholog of actin that has diverged to perform specialized roles in bacterial cell morphogenesis, cell wall organization, and growth.

In this study, we have demonstrated that bacterial actin homolog MreB, could be a potential target for neighbouring bacteria during interbacterial competition. Pathogenic bacteria often target the eukaryotic cytoskeletal protein actin directly or via actin regulators to establish infection. *Shigella* IcsA promotes actin-tail formation to drive intracellular and cell-to-cell movement, while *Salmonella* Typhimurium uses the T3SS effector SopE to trigger WAVE2-and Arp2/3-mediated actin rearrangement during epithelial invasion^65,66^. *V. cholerae* T6SS-component ACD_VC_ catalyzes an isopeptide bond between the γ-carboxyl group of Glu270 and the ε-amino group of Lys50, crosslinking actin monomers both *in vivo* and *in vitro*, causing cell rounding^33^. Beyond targeting mammalian host cells, pathogenic bacteria also compete with neighbouring bacterial cells for occupying the niche and nutrition acquisition. This competition is generally mediated via secretion of various toxins. Using MD simulation study, here we found *Vibrio cholerae* T6SS toxin ACD_VC_ made a stable complex with *E. coli* MreB (Figure 2, Video S1). This interaction was further validated experimentally by high-speed co-sedimentation and ELISA, demonstrating that ACD_VC_ directly binds MreB (Figure 3D, E). Notably, despite this robust interaction, ACD_VC_ failed to crosslink MreB, in sharp contrast to its well-established actin-crosslinking activity toward mammalian actin (Figure 4A). These findings reveal that ACD_VC_ might regulate MreB through a mechanism distinct from its canonical actin-crosslinking activity. Comparison of the crystal structure of rabbit actin with the high-confidence AlphaFold model of *E. coli* MreB had represented a plausible explanation of the above. Although MreB retains the conserved Lys27 (corresponds to K50 of actin) residue located within subdomain IB, it lacks the corresponding Glu270 residue in subdomain IIB that serves as the crosslinking partner in actin (Figure 4B). Structural alignment identified Glu260 of MreB as the closest positional equivalent of actin Glu270. But there is a striking difference in the spatial arrangement of these residues in MreB, compared to actin (Figure 4B, S2). Altered spatial organization of Lys27 and Glu260 in MreB is therefore likely to prevent formation of the intermolecular isopeptide bond catalyzed by ACD_VC_, explaining why MreB is not crosslinked despite sharing the conserved actin fold (Figure 4B, C, D). These observations indicate that conservation of individual residues alone is insufficient for ACD_VC_-mediated crosslinking. Instead, the precise three-dimensional arrangement and orientation of the reactive residues appear to be critical determinants of substrate recognition and catalysis.

Although the functional consequences of MreB-ACD_VC_ have not been deciphered yet, our findings are consistent with previous reports demonstrating that the bacterial cytoskeleton component, MreB, may serve as a crucial target for bacterial toxins during competition. For example, *Escherichia coli* toxins CbtA, CptA, and YeeV directly target the bacterial cytoskeletal proteins FtsZ and MreB, disrupting their polymerization and thereby impairing cell division. Apart from this, some of the T6SS effectors are also known to extend their effects towards bacterial cells for competitive fitness in polymicrobial communities. Like, the marine isolate *Vibrio anguillarum* uses T6SS mediated contact dependent antagonism to access nutrients from neighbouring cells. Whereas *Pseudomonas syringe* employs T6SS-1 effectors Tde1 and Tde4 to degrade *E. coli* DNA and cause changes in cell morphology. This competition is not confined to natural environments. In the mouse gut, *Salmonella enterica* serovar Typhimurium uses the T6SS effector Tae4 to kill commensal *Klebsiella oxytoca*, while in zebrafish, *V. cholerae* uses its T6SS to enhance intestinal contraction and expel the symbiont *Aeromonas veronii*, rather than killing it directly. Collectively, our findings establish, for the first time, that a T6SS effector *V. cholerae* ACD_VC_, targets the bacterial actin homolog MreB of *E. coli*, expanding its known target repertoire beyond mammalian actin.

We next asked whether this interaction nonetheless carries functional consequences at the cellular level. Expression of ACD_VC_ in *E. coli* cells at 40°C resulted in a pronounced growth inhibition, with ACD_VC_-expressing cells exhibiting an approximately 100-fold reduction in survivability compared with the control (Figure 5A). This observation was further corroborated by colony-forming unit (CFU) enumeration, which confirmed a statistically significant decrease in viable cells (Figure 5B). This suggested that, ACD_VC_-MreB interaction might hamper the cellular MreB dynamics leading to decrease in cell survivability. However, expression of ACD_VC_ at 37°C did not produce a significant inhibition in cell viability (Figure S6) The enhanced toxicity observed at elevated temperature may be associated with the temperature-dependent regulation of the *V. cholerae* Type VI secretion system (T6SS). Previous studies have shown that expression of the T6SS marker gene *hcp* is strongly influenced by temperature, remaining low at 15°C but increasing more than 50-fold at 25°C, indicating that T6SS-associated effectors are responsive to thermal stress^67^. The increased toxicity at 40°C suggests that elevated temperature may enhance ACD_VC_ activity, stability, or its interaction with MreB. Together, these results show that the MreB-ACD_VC_ interaction compromises bacterial viability, even though ACD_VC_ does not chemically crosslink its target, indicating that crosslinking is not the only route by which ACD_VC_ can exert toxicity. In addition to the reduction in cell viability, ACD_VC_ expression induced pronounced cell elongation accompanied by distinct lateral bulges, indicating that perturbation of MreB function alters bacterial cell morphology (Figure 6). MreB is the principal bacterial actin homolog responsible for maintaining rod-shaped morphology by coordinating cell wall synthesis during elongation^68,69^. Early genetic studies demonstrated that loss of MreB converts rod-shaped *E. coli* cells into spherical cells, highlighting its essential role in morphogenesis^69^. Several proteins that interfere with bacterial cytoskeletal dynamics produce characteristic morphological phenotypes. For example, the SOS-induced protein SulA inhibits cell division by binding FtsZ and blocking its GTPase activity, resulting in filamentous cells^70,71^. Likewise, the toxin YeeV disrupts the polymerization of both MreB and FtsZ, producing a distinctive lemon-shaped morphology due to simultaneous defects in cell elongation and cytokinesis^72^. Notably, the phenotype observed upon ACD_VC_ expression differs from either of these established cytoskeletal perturbations, suggesting that ACD_VC_ may affect MreB through a distinct mechanism (Figure 6A). Previous studies have shown that elevated MreB levels can promote cell elongation and, in some cases, increase cell width by altering cytoskeletal organization and delaying cell division^73^. This leads us to favor a model in which ACD_VC_ binding may stabilize MreB filaments or reduce their turnover, thereby disrupting the dynamic remodelling required for normal cell elongation and division. An alternative, but currently untested, possibility is that ACD_VC_ indirectly increases the cellular abundance of functional MreB. Future studies examining MreB polymer dynamics and protein abundance will be required to distinguish between these mechanisms. A further notable feature of the ACD_VC_ phenotype was the frequent positioning of lateral bulges near mid-cell, coinciding with the expected location of the division septum (Figure 6A, D). This spatial pattern might be suggestive in light of MreB’s known coordination with the FtsZ Z-ring during cytokinesis^74^. During bacterial cell division, MreB reorganizes from a transient singlet into a mature doublet ring that flanks the central FtsZ ring at the division septum. The MreB doublet positions itself on either side of FtsZ, helping maintain the division machinery at midcell^75^. This MreB ring also contains MreC, MreD, PBP2, and RodA. Formation of the ring requires FtsZ, while MreB, MreC, MreD, and RodA can assemble independently^76^. Collectively, the occurrence of multiple bulges on elongated cells strongly points towards defective cell division due to MreB-ACD_VC_ interaction. We therefore propose that ACD_VC_-mediated stabilization or dysregulation of MreB might causes its abnormal accumulation at sites of septal assembly, and elongation of the cell, producing localized defects that manifest as lateral bulges and cell filamentation (Figure 7). While this model will require direct experimental validation, it offers a plausible mechanistic explanation for the distinctive morphological phenotype seen in ACD_VC_-expressing cells.

**Figure 7:**
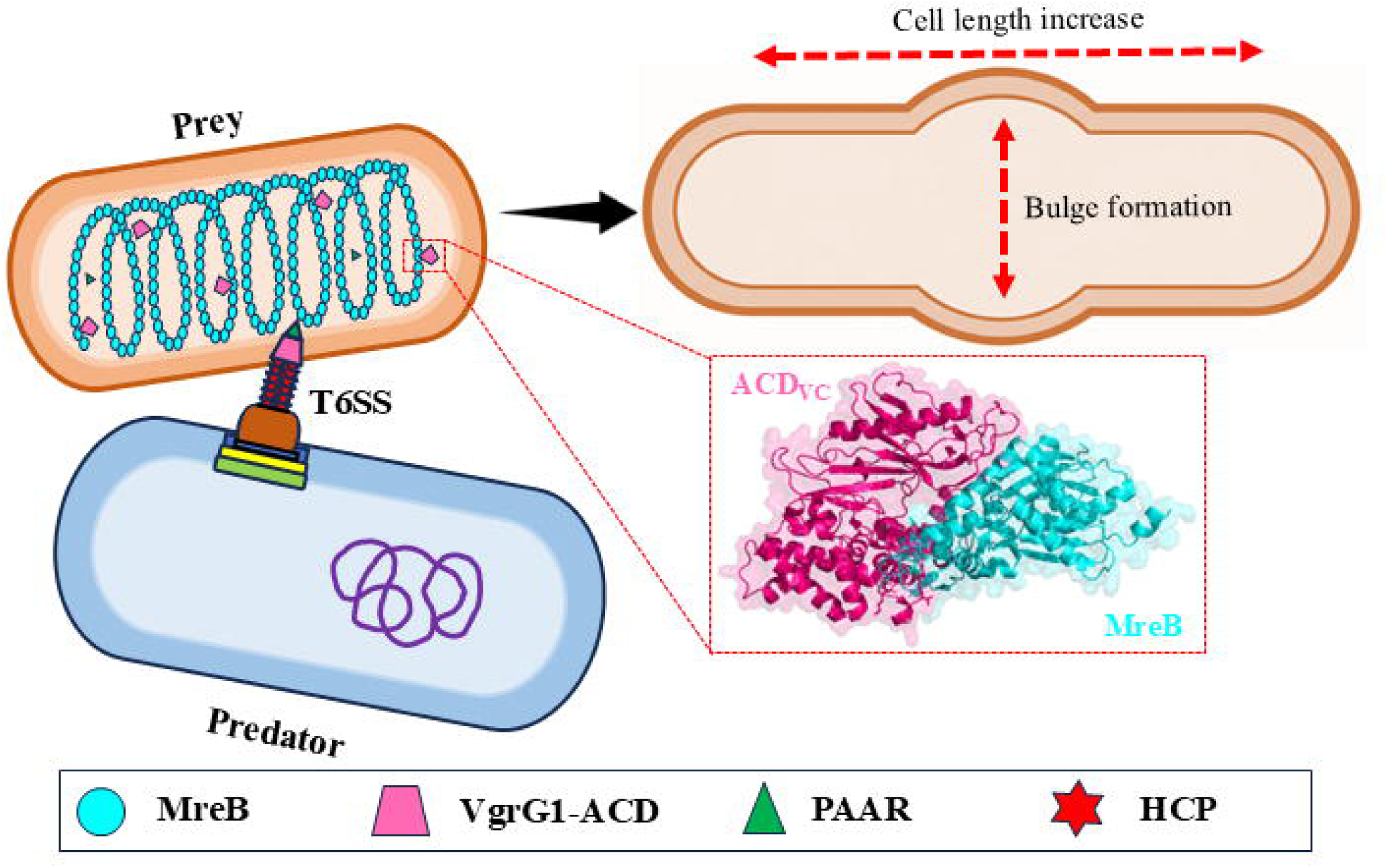
T6SS effector ACD_VC_ targeting bacterial cytoskeleton MreB to induce morphological alterations. The predator bacterium utilizes its T6SS machinery to inject the VgrG1 C-terminal Actin-Crosslinking Domain (ACD_VC_) into the target prey cell. Translocated ACD_VC_ directly binds to the host cytoskeletal protein MreB (cyan; inset), to alter its dynamics and results into cell length extension and bulge formation. Structural components of the T6SS spike and tube assembly-including PAAR (green triangle), Hcp (red star), and VgrG1-ACD (pink diamond)-are indicated in the box.

Collectively, our findings uncover a previously unrecognized cross-kingdom targeting strategy in which the *V. cholerae* T6SS effector ACD_VC_ exploits distinct molecular mechanisms to antagonize mammalian actin and the bacterial actin homolog MreB. This mechanistic versatility broadens the functional repertoire of ACD_VC_, provides a new framework for understanding T6SS-mediated interbacterial competition, and offers important insight into how *V. cholerae* may establish a competitive advantage within complex polymicrobial gut ecosystems.

## 4. Materials and Methods

### 4.1. Multiple sequence alignment and Phylogenetic tree construction

Protein sequences of representative actin superfamily members, including mammalian actin and actin homologs from archaea, bacteria, plants, and yeast, were retrieved from the NCBI database (Table S4). For phylogenetic tree construction, protein sequences were aligned using MUSCLE in MEGA11 software^77^. Then, large gaps and poorly aligned tails were trimmed manually. Tree was constructed using Maximum Likelihood method with Gamma-distributed rate variation among sited (G) and 1000 bootstrap replicates. Generated tree was visualized using iTOL (Interactive Tree of Life) v7 software^78^.

Percentage identity matrix was generated after Multiple sequence alignment (MSA) of protein sequences using Clustal Omega and values were plotted in GraphPad Prism v8.4.2.

### 4.2. Structure prediction, Molecular docking and Simulations

The amino acid sequences of *Escherichia coli* MreB (UniProt ID: P0A9X4) and the Actin Crosslinking Domain (ACD_VC_; residues 732-1163) of VgrG1 from *Vibrio cholerae* O139 (UniProt ID: Q9KS45) were retrieved from UniProt. The three-dimensional (3D) structures of MreB, ACD_VC_ other proteins were predicted individually using AlphaFold2 (ColabFold) with the AlphaFold2_ptm model optimized for monomeric protein prediction^79^. The quality of the predicted models was evaluated using the predicted aligned error (PAE) and per-residue predicted local distance difference test (pLDDT) scores generated by the server.

Protein-protein docking was performed using the AlphaFold2_multimer_v3 model to predict the MreB-ACD_VC_ complex^80^. The highest-confidence predicted complex was selected for further analysis. The resulting Protein Data Bank (PDB) file was visualized in PyMOL to identify the interacting interface between the two proteins^81^. Amino acid residues located within 2.5 Å of each other were considered potential interacting residues.

The structural stability of the predicted MreB-ACD_VC_ complex was further evaluated by molecular dynamics (MD) simulations using GROMACS (version 2024.4)^55^. The complex was solvated in a cubic simulation box containing SPC216 water molecules, and counterions were added to neutralize the system. Following energy minimization to eliminate steric clashes and obtain a stable starting conformation, the system was equilibrated before a 50 ns production MD simulation was performed under periodic boundary conditions. The stability and compactness of the complex were assessed by calculating the root mean square deviation (RMSD) and radius of gyration (Rg) throughout the simulation. Structural dynamics and conformational changes of the MreB-ACD_VC_ complex were further analyzed by visualizing the simulation trajectories (.xtc files) in PyMOL^82^.

### 4.3. DNA isolation, Cloning and Plasmid Construction

*E. coli* DH5α genomic DNA was isolated by CTAB method^83^. The *mreB* gene (Uniprot ID: P0A9X4, amino acids 1-347) was amplified and subsequently cloned into 6X-His tagged pET28a(+) vector (Novagene)^36^. ACD_VC_ construct (amino acids 732-1163), used in this study, had obtained from our previous study^35^.

### 4.4. Protein expression and Purification

Constructs were transformed into *E. coli* BL21 (DE3) cells and cultured at 37°C with 30 µg/mL kanamycin, until OD_600_ reached 0.5. Protein expression was induced by 0.5 mM IPTG at either 18°C (for ACD_VC_) or 37°C (for MreB) for 12 hours and 4 hours, respectively. Cells were harvested by centrifugation at 4000 rpm for 10 mins^84^.

For 6X-His-tagged protein purification cells were resuspended in lysis buffer containing 50mM Tris pH 8, 100 mM NaCl, 30 mM Imidazole, 0.2% IGEPAL, 1X PIC (phenylmethylsulfonyl fluoride, benzamidine hydrochloride, leupeptin, aprotinin, pepstatin A), 0.5 mM DTT and, 5% Glycerol. For MreB purification, 0.2 mM ATP, mM MgCl_2,_ 0.5 mg/mL lysozyme were added in lysis buffer. The cell suspension was then sonicated for 5 mins. The lysates were centrifuged at 12000 rpm for 10 mins. Supernatants were collected and incubated with Ni-NTA agarose beads for 2 hours at 5 rpm rotation at 4°C. Beads were collected by centrifugation at 1500 rpm for 2 mins and washed thrice by wash buffer containing 50 mM Tris pH 8, 150 mM NaCl and, 30 mM Imidazole. Proteins were eluted using elution buffer (50 mM Tris pH 8, 100 mM NaCl and, 350 mM Imidazole and, 5% Glycerol). Purified proteins were dialysed against dialysis buffer containing 5 mM Tris pH 7.5, 0.1 mM CaCl_2_, and 0.2 mM ATP^56,85^.

### 4.5. High-speed co-sedimentation assay

Since MreB, like eukaryotic actin, undergoes ATP-dependent polymerization into filamentous structures, establishing appropriate polymerization conditions was essential before investigating its interaction with ACD_VC_. However, the biochemical requirements for MreB polymerization differ considerably among bacterial species. For instance, *Thermotoga maritima* MreB polymerizes in KMEI buffer containing 0.2 mM ATP, 10mM imidazole pH 7.0, 20 mM KCl, 1mM MgCl_2_ and 1mM EGTA^86^, whereas *Bacillus subtilis* MreB has been reported to polymerize optimally in 5 mM Tris-HCl, 0.1 mM CaCl_2_, 0.2 mM ATP, pH 7.5^87^. Therefore, polymerization conditions for *E. coli* MreB were optimized to subsequent interaction studies with ACD_VC_.

In this study, *E. coli* MreB was polymerized under low-salt buffer conditions containing 5 mM Tris-HCl (pH 7.5), 0.1 mM CaClL, and 0.2 mM ATP. Polymerization was initiated by the addition of 5 mM MgClL, followed by incubation at 25°C for 30 min^58^. The resulting filaments were sedimented by ultracentrifugation at 90,000 rpm for 30 min, with centrifugation performed at either 25°C or 4°C to evaluate filament stability. When centrifugation was carried out at 25°C, MreB was recovered in the pellet fraction, indicating efficient filament formation and stability under these conditions (Figure S4). In contrast, centrifugation at 4°C resulted in a substantial proportion of MreB remaining in the supernatant, with nearly half of the protein failing to sediment (Figure S4). Based on these results, subsequent MreB polymerization and co-sedimentation assays were performed with ultracentrifugation at 25°C to preserve filament integrity. For the test samples, MreB was added with increasing concentrations of ACD_VC_ before the addition of 5 mM MgClL. The reaction mixtures were then incubated at 25°C for 30 min. The reaction was centrifuged at 90000 rpm at 25°C for 30 mins. The supernatant fraction was collected and the pellet fraction was resuspended with an equal volume of buffer. The sample was mixed with sample loading buffer and run in 10% SDS-PAGE^88^.

MreB was used as a positive control to verify sedimentation following high-speed centrifugation. In contrast, the protein control, ACD_VC_, was used to confirm protein solubility, as it remained in the supernatant fraction even after centrifugation.

### 4.6. Antibody

Polyclonal antisera against MreB were generated by immunizing BALB/c mice with purified His-tagged *Escherichia coli* MreB. The immunization protocol was carried out over 70 days following approval from the Institutional Animal Ethics Committee (IAEC), under protocol number IISER/IAEC/AP/2024/138. Terminal bleeds were collected, and the specificity of the antisera was validated by western blot analysis using purified recombinant His-tagged MreB^89,90^.

### 4.7. Enzyme-linked immunosorbent assay (ELISA)

To validate the direct interaction between MreB and ACD_VC_ we performed ELISA. Purified ACD_VC_ (amino acids 732 - 1163) was coated onto flat bottom ELISA plate (MaxiSorp^TM^), in a concentration of 10 µg in each well and 1X PBS was taken as control. Plate was incubated overnight at 4°C and remaining steps were done in room-temperature. Blocking was with 5% BSA, prepared in 1X PBS for 2 hours, followed by incubation of MreB, prepared in 1X PBS, in a concentration-dependent manner (5, 2.5, 1.25, 0.625, 0.312, 0.156, 0.078 µg/mL) for 2 hours. The interaction was detected by three steps, (1) addition of mice raised anti-MreB primary antibody (1:1000 dilutions) for 2 hours, (2) incubation with HRP-conjugated goat raised anti-mice IgG secondary antibody (1:50000 dilutions, Invitrogen) for 45 mins, (3) addition of 1X Tetramethylbenzidine (TMB, Sigma Aldrich) for 10 mins to develop the colour. The reaction was stopped by adding 5N H_2_SO_4_. After each step, washing was done thrice using 1X PBST (0.02% Tween-20 in 1X PBS). Finally, the absorbance was measured using a microplate reader (Epoch2) at 450 nm^89^.

### 4.8. MreB crosslinking assay

Purified MreB was incubated with various concentrations of ACD_VC_ in polymerisation buffer (5 mM Tris-HCl (pH 7.5), 0.1 mM CaClL, 0.2 mM ATP, and 5 mM MgCl_2_) at room temperature for 45 mins. Whereas Rabbit Muscle Actin (RMA) was incubated with ACD_VC_ in the same buffer and served as a positive control. Samples were then mixed with sample loading buffer and run in 10% SDS-PAGE^36^.

### 4.9. Cell survivability assay

To check the effect of ACD_VC_ on cell survivability, we had transformed 6X-His tagged ACD_VC_ construct into *E. coli* BL21 (DE3) cells. As a negative control we have taken pET28a(+) vector and 6X-His tagged mouse Profilin (mPfn) and also transformed into BL21 (DE3) cells. For primary culture, Cells were grown in LB media, supplemented with 30 mg/mL kanamycin overnight at 37°C. Next day, 1% culture was used for giving secondary culture supplemented with 30 mg/mL kanamycin at 37°C until OD_600_ reached at 0.5. Cells were induced with 0.5 mM IPTG at either 40°C and 37°C for 4 hours. Each bacterial culture was serially diluted upto 10^-10^. 5 µl of samples from each dilution was then spotted on LB agar plates, containing 30 mg/mL kanamycin and incubated for overnight at 37°C.

For CFU count, 40 µl of diluted samples from different dilutions were spread onto kanamycin supplemented LB agar plates.

### 4.10. SEM imaging and Bacterial length measurement

To analyze the effect ACD_VC_ in regulating cell morphology, we did SEM imaging followed by measurement of bacterial cell length and width. Cells were grown and induced in LB media as previously described method.

For SEM sample preparation, 1 mL culture from each flask was taken in 1.5 mL in microcentrifuge tube. Cells were then collected by centrifugation at 5000 rpm from 5 mins. Supernatant was discarded and the pellet was washed twice with filtered autoclaved Milli-Q water, followed by washing thrice with 1X PBS. After each wash cells were pelleted down by centrifugation at 5000 rpm for 5 mins. Cells were fixed with 400 µL of 2.5% glutaraldehyde (prepared in 1X PBS) for 30 mins in dark. Then centrifugation was done and supernatant was discarded. Pellet was washed with filtered autoclaved Milli-Q water once and 1X PBS trice. After centrifugation pellet was resuspended in 1% osmium tetroxide (OsO_4,_ prepared in 1X PBS) for 1 hour in dark. The pellet was washed again in water thrice. Cells were dehydrated through a series of ethanol concentration gradient: 10%, 30%, 50%, 70%, 90%, 100%. During dehydration, cells were resuspended and kept in specific ethanol concentrations with mild agitation for 10 mins. With 100% ethanol cells were kept for 1 hour with no agitation. Then, ells were centrifuged once with water and washed thrice with 1X PBS. Finally, cells were diluted in 1 mL of 1X PBS and drop-cast on 12 mm cleaned coverslip, followed by air dry. All the steps were done at room temperature.

All images were taken using either FESEM SUPRA 55 VP (CARL ZEISS) or FESEM JSM-IT800 (JEOL) at 5kV. Images were processed and labelled using ImageJ software. For length and width measurement, 33 cells were considered from different fields and from three independent experiments for each sample.

### 4.11. Statistical analysis

Data were plotted and analysed statistically using One-way ANOVA with post hoc Tukey HSD test (*P* < 0.05), respectively using GraphPad Prism (v8.4.2.) software.

## Abbreviations

T6SS: Type VI secretion system
Tss: Type six secretion
VgrG1: Valine-glycine repeat protein G1
ACD: Actin cross-linking domain
HCP: Hemolysin coregulated protein
CTX: Cholera toxin

## Data availability

All supporting data of this manuscript and findings are available to corresponding author upon valid request.

## Acknowledgement

The authors thanks to IISER Kolkata and Department of Biological Sciences for funding and facilities. Amaresh Jana acknowledges IISER Kolkata for providing fellowship. Amrita Maity acknowledges Department of Biotechnology (DBT) for providing fellowship and Dr. Dipjyoti Das for insightful comments. We thank IISER Kolkata Central Instrument Facility for providing FESEM instrument, and the Department of Biological Sciences for the computational facility.

## Author contributions

**Amaresh Jana** and **Amrita Maity:** Experimentation; data curation; visualization; validation; methodology; writing-original draft; review and editing. **Shubham Das:** Investigation; visualisation; review and editing **Saikat Das:** Visualisation; writing-original draft. **Priyanka Dutta:** Validation; review and editing **Sankar Maiti:** Project administration; investigation; conceptualization; funding acquisition; writing-original draft; review and editing.

## Conflict of interest

Authors declare having no conflict of interest.

## Notes

### Competing Interest Statement

The authors have declared no competing interest.

