## Supplementary file for "*Vibrio cholerae* VgrG1-ACD Targets *E. coli* MreB and Regulates Cellular Morphology"

**This file includes:**

Table S1 to S4

Figure S1 to S7

Video S1

**Table S1: List of primers used in this study**

| Name of constructs | Vector backbone | Primer sequences | Source |
|---|---|---|---|
| MreB | pET28a(+) | Forward primer<br>5'-CGCGCGGGATCCATGTTGAAAAAATTTCGTGGC-3'<br>Reverse primer<br>5'-GCGCGCCTCGAGTTACTCTTCGCTGAACAGGTC-3' | This paper |
| VgrG1 ACD (732-1163) | pET28a(+) | Forward primer<br>5'-GGAAGATCTCCAACACATTTCCCGAAGTC-3'<br>Reverse primer<br>5'-CGCAAGCTTTTATTGCCATTCTTGAGGATTATGC-3' | Dutta *et al.* 2019 |

**Table S2: List of reagents used in this study**

| | Source | Cat. no |
|---|---|---|
| **Bacterial competent cells** | | |
| XL10 Gold | Stratagene, Agilent Technologies | 200314 |
| BL21(DE3) | Agilent Technologies | 200131 |
| **Recombinant DNA** | | |
| pET28a(+) | Novagen, Merck Life Science | 69864 |
| **Chemical, Enzymes and other Reagents** | | |
| BamHI | NEB | R0136S |
| XhoI | NEB | R0146S |
| T4 Ligase | Thermo | EL0011 |
| IGEPAL | Merck | 56741 |
| EDTA | Sigma Aldrich | E4884 |
| Imidazole | US Biologicals | 452259 |
| DTT | HiMedia | MB070 |
| EGTA | VWR | 20308.156 |
| HEPES | Biowest | P5455 |
| Glycerol | Sigma-Aldrich | G7893 |
| ATP | Abcam | AB156525 |
| MgCl2 | Sigma-Aldrich | M8266 |
| IPTG | HiMedia | RM2578 |
| Ni-NTA Beads | Qiagen | 30210 |
| Glacial acetic acid | Merck | 1.93402 |
| Potassium chloride | Merck | P3911 |
| Trizmabase | Merck | 93362 |
| Sodium chloride | Merck | S9888 |
| Glutaraldehyde | Merck | G5882 |
| Ammonium persulfate | Merck | 248614 |
| TEMED | Merck | T9281 |
| Acetone | Merck | SI9F690655 |
| Methanol | FINAR | 389302C250 |
| Sodium dodecyl sulfate | Sigma-Aldrich | L4509 |
| Protease inhibitor cocktail | Sigma-Aldrich | P8340 |

**Table S3: Software used in this study**

| **Softwares used** | | |
|---|---|---|
| UniProt | https://www.uniprot.org/ | N/A |
| PyMOL (v2.5) | https://pymol.org/ | N/A |
| MEGA | https://www.megasoftware.net/ | N/A |
| GROMACS (v2021) | https://www.gromacs.org/ | N/A |
| Clustal Omega | https://www.ebi.ac.uk/jdispatcher/msa/clustalo | N/A |
| Fiji ImageJ.2 | https://imagej.net/software/fiji/ | N/A |
| iTOL | https://itol.embl.de/ | N/A |
| GraphPad Prism 8 | https://www.graphpad.com/ | N/A |

**Table S4: Protein used of phylogenetic tree construction and structural comparison**

| **Protein** | **Organism** | **Accession no.** |
|---|---|---|
| MreB | E. coli | AAA83891.1 |
| MreB | B. subtilis | AAA22605.1 |
| MreB | H. influenzae | AAC21715.1 |
| MreB | S. typhimurium | AAL22243.1 |
| MreB | S. flexneri | AAN44753.2 |
| MreB | T. maritima | AAD35673.1 |
| MreB | V. cholerae | AAF93588.1 |
| MreB | P. aeruginosa | AAG07869.1 |
| MreB | B. thailandensis | ABC39383.1 |
| MreB | K. pneumoniae | CDL08726.1 |
| MreB | H. pylori | AAD08416.1 |
| ParM | E. coli | CAA31264.1 |
| Mbl | B. subtilis | AAA67878.1 |
| MreBH | B. subtilis | BAA07047.1 |
| AlfA | B. subtilis | WP_013603336 |
| FtsA | E. coli | AAA23817.1 |
| FtsA | B. subtilis | AAA22456.1 |
| FtsA | S. aureus | AAC45628.1 |
| FtsA | P. aeruginosa | AAA95992.2 |
| Alp7A | B. subtilis | ACU27363.1 |
| Alp7A | P. megaterium | AJI25597.1 |
| MamK | P. magneticum | BAE49769.1 |
| MamK | M. gryphiswaldense | CAE12034.1 |
| Salactin | H. salinarum | AAG18772.1 |
| Volactin | H. volcanii | ADE04559.1 |
| Crenactin | P. calidifontis | ABO09052.1 |
| Actin | T. adornatum | AGT35143.1 |
| Actin | H. sapiens | AAB59376.1 |
| Actin | O. cuniculus | CAA24241.1 |
| Actin | S. cerevisiae | CAA24597.1 |

Figure S1

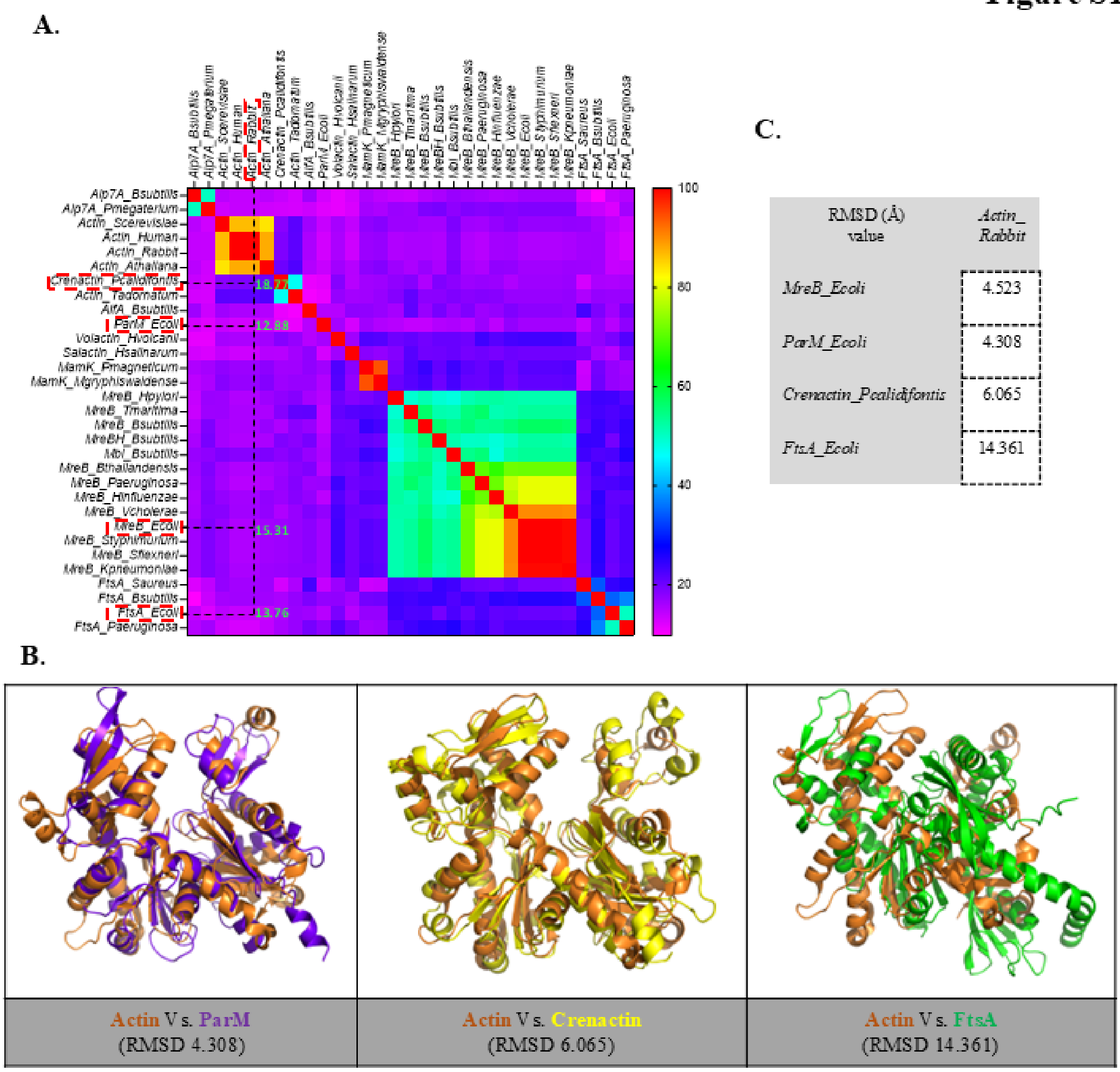


| RMSD (Å) value | *Actin_ Rabbit* |
|---|---|
| *MreB_Ecoli* | 4.523 |
| *ParM_Ecoli* | 4.308 |
| *Crenactin_Pcalidifontis* | 6.065 |
| *FtsA_Ecoli* | 14.361 |

**Figure S1: Sequence and structural comparison of representative actin superfamily proteins. (A)** Heat-map representing the sequence percentage identity among selected actin family proteins including, rabbit actin, crenactin, MreB, FtsA, ParM and other actin homologs. Color shading (Violet to Red) represents gradual increase in sequence identity. **(B)** Structural superposition of *E. coli* ParM (Blue; PDB: 1MWM), *P. calidifontis* Crenactin (Yellow; PDB: 5LY3) and *E. coli* FtsA (Green; PDB: 7Q6D) with actin (Yellow; PDB: 1J6Z). **(C)** RMSD value of superposition structures from bacterial actin homolog with rabbit actin.

#### Figure S2

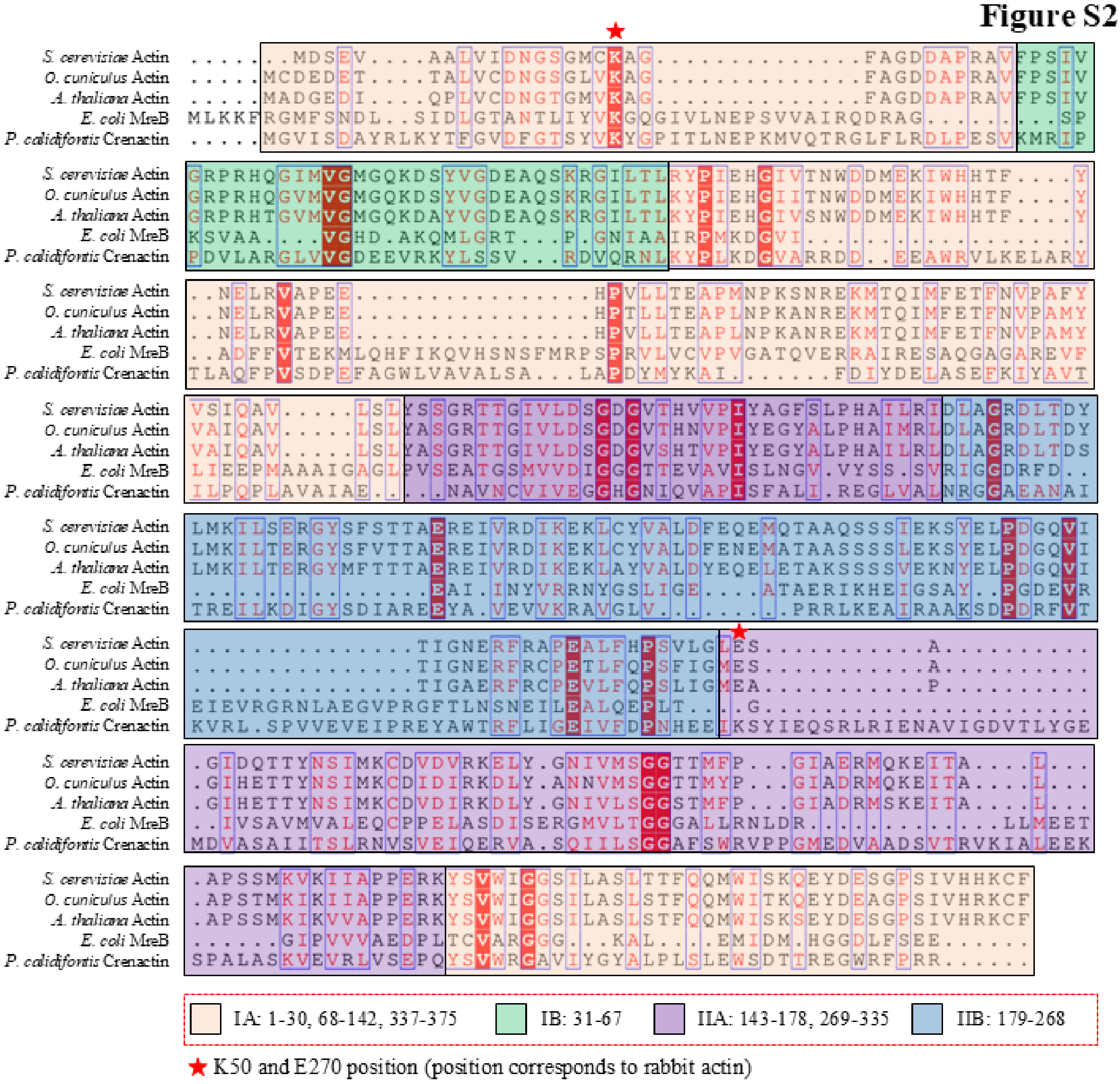


**Figure S2**: **Sequence conservation and domain organisation of MreB with eukaryotic Actins.** Multiple Sequence Alignment (MSA) result showed preservation of core amino acids in each sub-domain (IA, IB, IIA, and IIB) between *E. coli* MreB and other eukaryotic actin homologs despite of sequence divergence. MSA color scheme: red background, identical residues; red letters, conserved residues; blue frame, highly similar residues; black letters, low conservation positions. Four actin-fold sub-(domains and the respective amino acid positions are highlighted in box below: IA: 1-30, 68-142, 337-375, IB: 31-67, IIA: 143-178, 269-335, IIB: 179-268. Red star mark represents K50 and E270 position (corresponds to rabbit actin)

### Figure S3

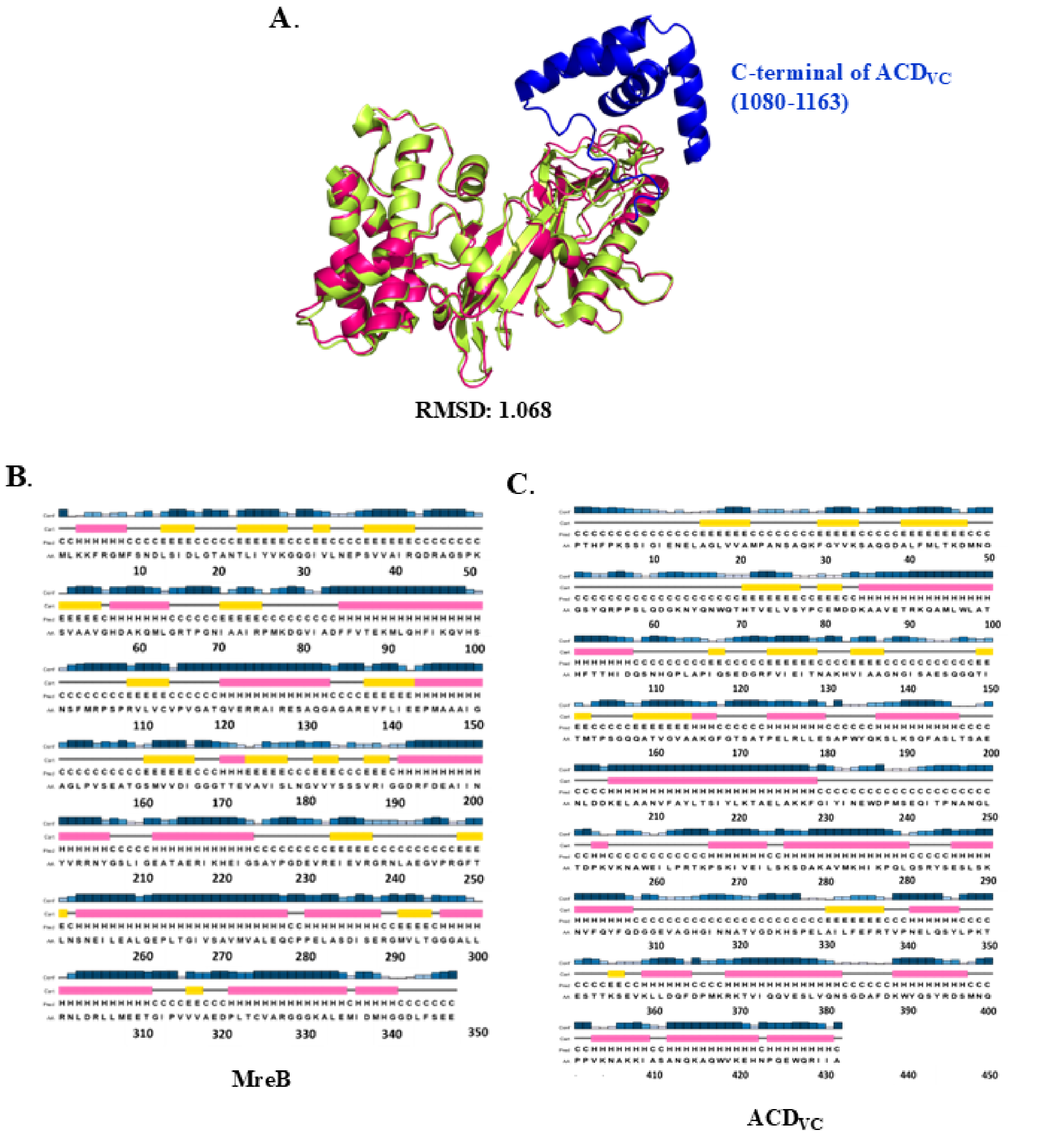


**Figure S3: Structural validation and secondary structure prediction of ACD$_{VC}$ and MreB.** **(A)** Structural alignment of experimentally resolved ACD$_{VC}$ (Green, PDB: 4DTD) and AlphaFold predicted model (Pink, c-terminal: blue) showed a RMSD of 1.068. Secondary structures of **(B)** MreB and **(C)** VgrG1 ACDvc was predicted through PSIPRED tool where yellow box represents beta sheet, pink box represent helix and grey line represents coil.

Figure S4

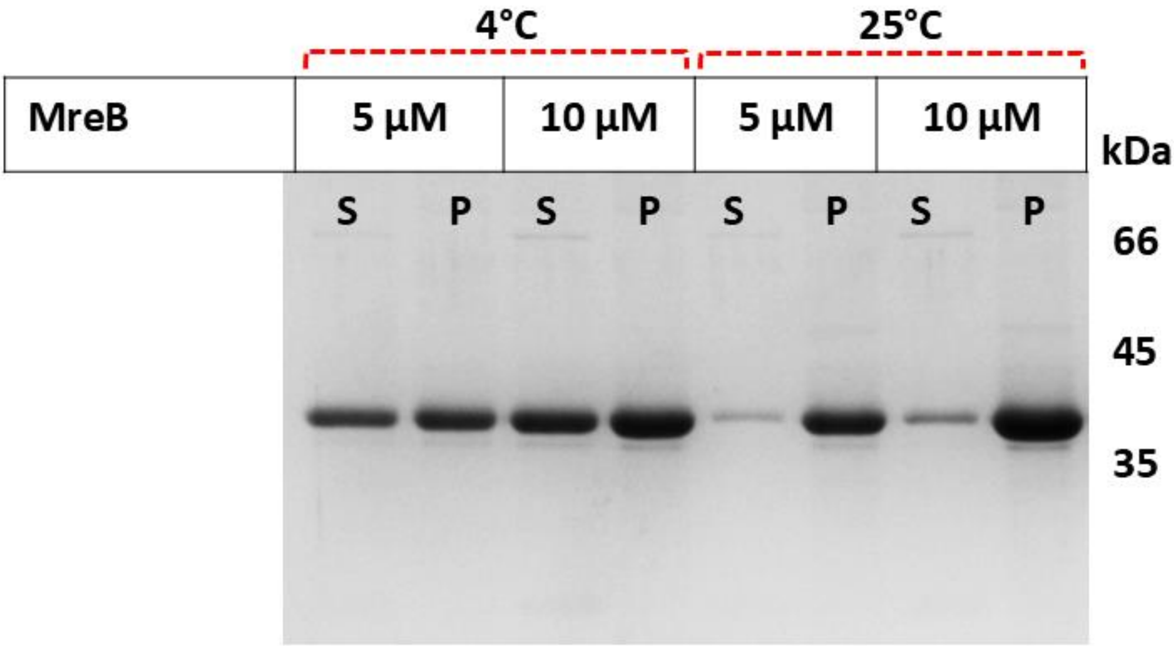


**Figure S4: At 4°C temperature MreB filament depolymerises.** Polymerised MreB fractions were centrifuged at 4°C and 25°C temperature. 10% SDS-PAGE shows, approximately half of MreB protein remains in supernatant fraction when centrifuged at 4°C. On the other hand, most of the protein went to the supernatant fraction when centrifuged at 25°C.

**Figure S5**

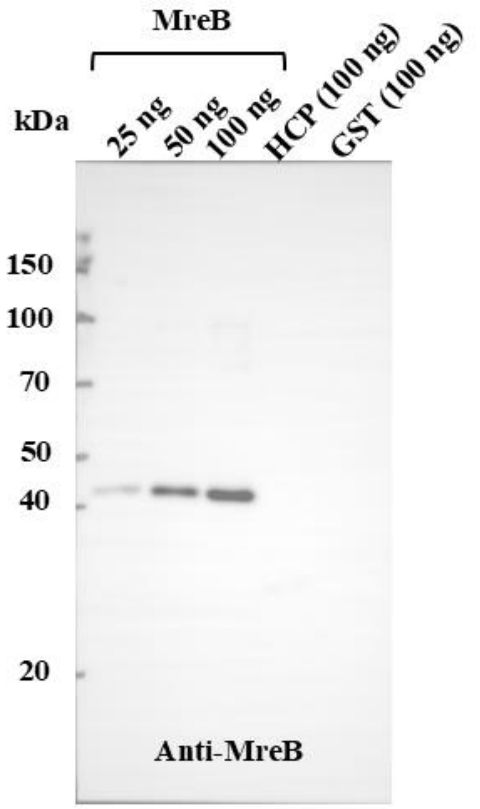


**Figure S5: Validation of anti-MreB polyclonal antibody specificity by immunoblotting.** Polyclonal antibodies were generated in mice against purified His-tagged *E. coli* MreB, and terminal bleed sera were evaluated by immunoblot analysis. Purified His-tagged MreB (25, 50, and 100 ng) was specifically detected as a single band at the expected molecular mass (~40 kDa) using the anti-MreB antisera (primary antibody, 1:1000; HRP-conjugated secondary antibody, 1:40,000). Purified His-tagged HCP and GST (100 ng each) were included as negative controls.

**Figure S6**

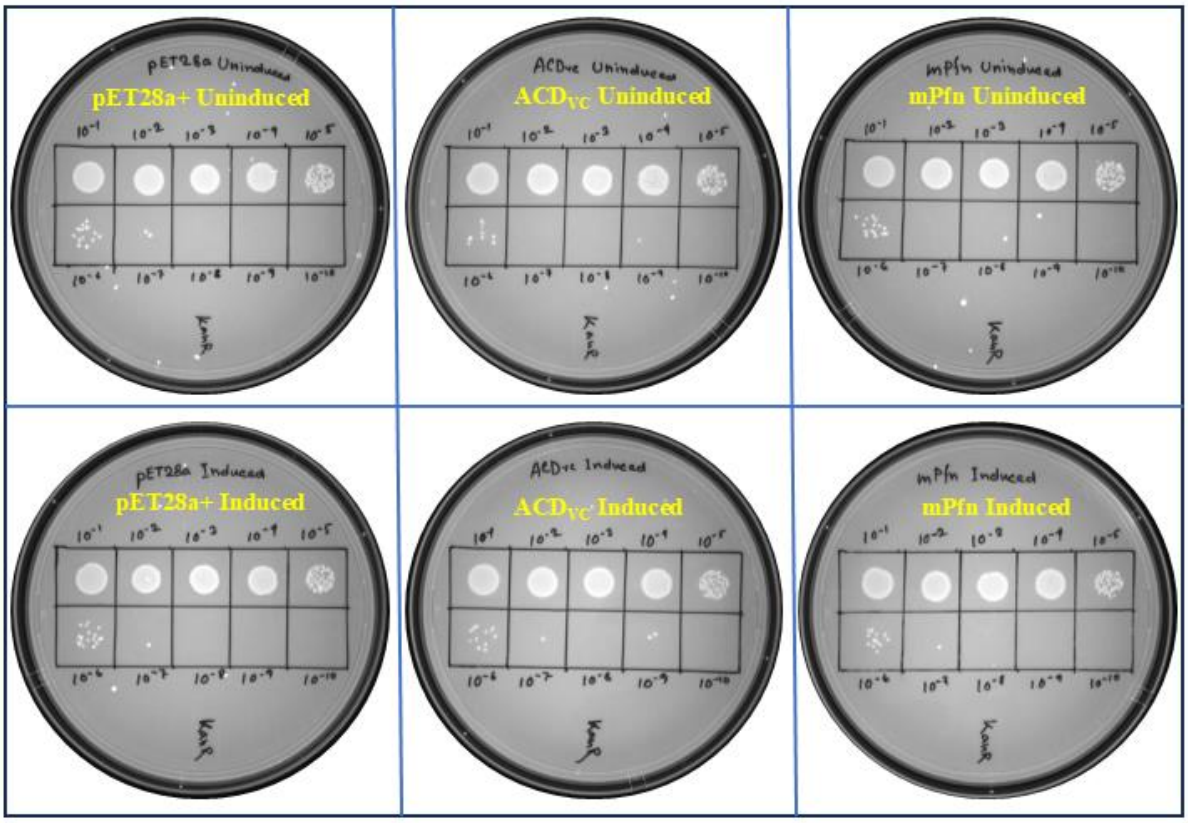


**Figure S6: ACD$_{VC}$ expression insignificant effect on *E. coli* cell survivability at 37°C.** Representative spot dilution assay showing the viability of *E. coli* BL21(DE3) cells harbouring pET28a(+), ACD$_{VC}$, and mPfn under uninduced and IPTG-induced conditions at 37°C. Cultures were serially diluted 10-fold ($10^{-1}$-$10^{-10}$), and 5 µL of each dilution was spotted onto LB-kanamycin agar plates. Data are representative of three independent experimental replicates (n = 3).

Figure S7

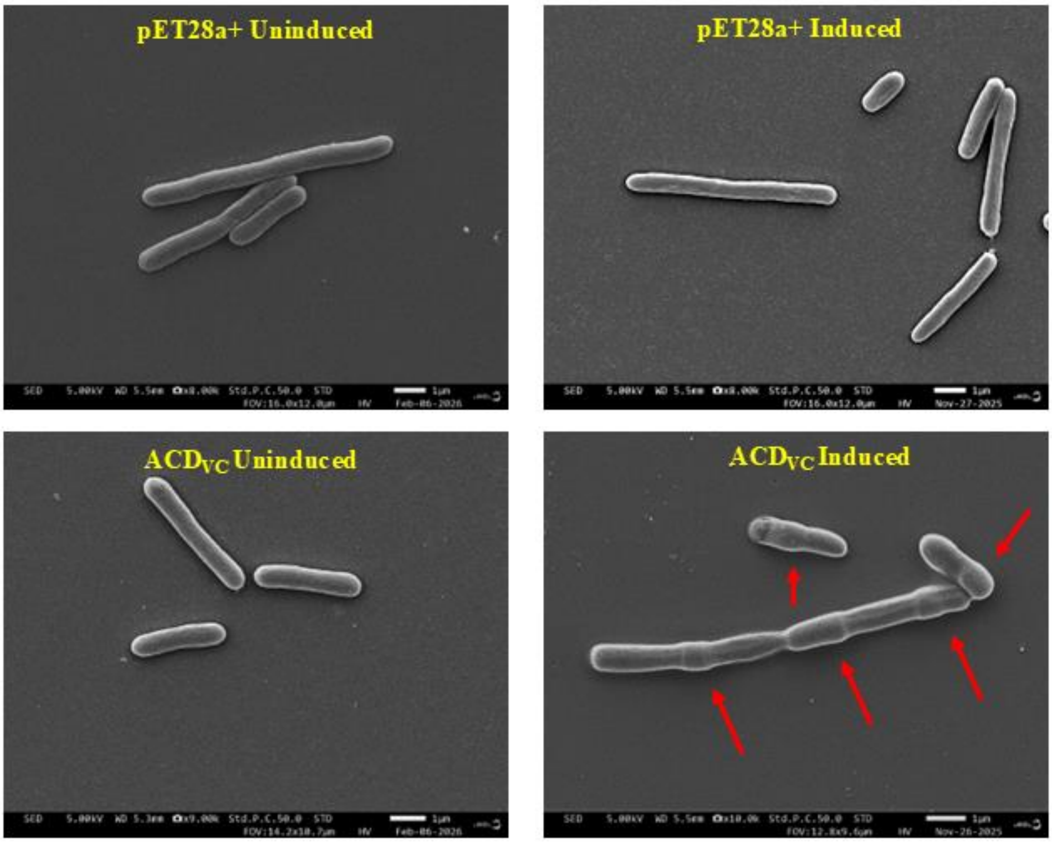


**Figure S7: ACD$_{VC}$ expression induces morphological abnormalities in *E. coli* at 37°C. (A)** Representative scanning electron microscopy (SEM) images of *E. coli* BL21(DE3) cells harbouring pET28a(+), and ACD$_{VC}$ under uninduced and IPTG-induced conditions at 37°C. Lateral bulges along the cell envelope in ACD$_{VC}$ induced cells are marked in red arrow. Images are representative of four independent biological experiments (n = 4). Scale bar, 1 µm.
